# Exploring Mobile Genetic Elements in *Vibrio cholerae*

**DOI:** 10.1101/2024.07.25.605194

**Authors:** Natália C. Drebes Dörr, Alexandre Lemopoulos, Melanie Blokesch

**Affiliations:** Laboratory of Molecular Microbiology, Global Health Institute, School of Life Sciences, Ecole Polytechnique Fédérale de Lausanne (EPFL), CH-1015 Lausanne, Switzerland

## Abstract

Members of the bacterial species *Vibrio cholerae* are known both as prominent constituents of marine environments and as the causative agents of cholera, a severe diarrheal disease. While strains responsible for cholera have been extensively studied over the past century, less is known about their environmental counterparts, despite their contributions to the species’ pangenome. This study analyzed the genome compositions of 46 *V. cholerae* strains, encompassing both pandemic and non-pandemic, toxigenic, and environmental variants, to explore the diversity of mobile genetic elements (MGEs) and bacterial defense systems. Our findings include both conserved and novel MGEs across the strains, pointing to shared evolutionary pathways and ecological niches. The defensome analysis revealed a wide array of antiviral mechanisms, extending well beyond the traditional CRISPR-Cas and restriction-modification systems. This underscores the dynamic arms race between *V. cholerae* and MGEs and/or bacteriophages and suggests that non-pandemic strains may act as reservoirs for emerging defense strategies. Moreover, the study showed that MGEs are integrated into genomic hotspots, which may serve as critical platforms for the exchange of defense systems, thereby enhancing *V. cholerae’s* adaptive capabilities against phage attacks and invading MGEs. Overall, this research offers foundational insights into *V. cholerae’s* genetic complexity and adaptive strategies, with implications for understanding the differences between environmental strains and their pandemic relatives, and the potential evolutionary paths that led to the emergence of pandemic strains.

## INTRODUCTION

Classifying prokaryotic genomes into specific units, such as species, is often a complex task due to their inherent genomic fluidity [1–4]. Comparative studies have shown that genomic variation within the same prokaryotic species is less about sequence variability and more about differences in gene content [1, 5]. These gene content variations are largely driven by mobile genetic elements (MGEs), which play a crucial role in bacterial evolution [6]. Indeed, the mobilome, representing the entire set of MGEs within a genome, is a significant component of the prokaryotic genome landscape. Although some MGEs can change their chromosomal positions within the same cell, they are most commonly transferred between different organisms’ genomes via horizontal gene transfer (HGT). This transfer typically occurs through mechanisms like transformation, conjugation, or transduction [3, 4, 6, 7].

The significance of MGEs in microbial evolution was first recognized through their role in pathogenicity [1, 8, 9]. Originally identified as large genetic elements unique to pathogenic isolates, carrying genes encoding virulence factors, they were termed “pathogenicity islands”. The acquisition of these elements enabled bacteria to transition from an environmental to a pathogenic lifestyle [1, 3, 9–12]. Subsequent research revealed that such islands are not exclusively associated with pathogens and can contain genes unrelated to virulence. Consequently, they are now referred to by the more encompassing term “genomic islands” [8, 12]. Indeed, the genes within such islands can contribute to various adaptive bacterial traits beyond pathogenicity, including the degradation of specific compounds, niche adaptation, symbiosis, interbacterial competition, defense against foreign DNA and phages, among others [3, 8, 12]. Genomic islands typically display unique features in contrast to the surrounding genome, such as an unusual GC content, being bordered by repeat sequences, and their common position adjacent to tRNA genes or other highly conserved regions of the genome. Despite their chromosomal integration, these islands often carry integrase-, transposase-, or conjugation-related genes, making them capable of mobilization [6, 8, 12].

*Vibrio cholerae* serves as an exemplary model to study the evolutionary impact of MGEs. This gram-negative bacterium is the causative agent of cholera, a severe diarrheal disease. However, most *V. cholerae* strains, and especially environmental isolates, are not implicated in the disease cholera. Notably, out of the more than 200 *V. cholerae* serogroups, only two (O1 and O139) are associated with historical and ongoing cholera pandemics [13–16].

Horizontally acquired MGEs have contributed to the classification of lineages from past and current cholera pandemics and allow us to distinguish distinct waves within the ongoing seventh pandemic [17–20]. Briefly, cholera-causing *V. cholerae* strains are characterized by the presence of *Vibrio* pathogenicity island 1 (VPI-1, also known as TCP island), which encodes the toxin-coregulated pilus (TCP). TCP is vital for intestinal colonization [21, 22] and serves as the receptor for the CTX bacteriophage (CTXΦ) [23]. The CTXΦ prophage harbors genes for the production of cholera toxin (CTX), the primary virulence factor responsible for the hallmark symptom of acute diarrhea [23]. In addition to these primary and virulence-encoding MGEs, cholera-causing strains also contain VPI-2. This island includes the *nan-nag* region, which plays a role in sialic acid metabolism [24]. Additionally, VPI-2 contains a cluster of phage-related genes and operons that encode established (DdmDE) or predicted phage and DNA defense systems [19, 24–28]. Moreover, pandemic strains responsible for the current 7^th^ cholera pandemic, often referred to as 7^th^ pandemic El Tor strains or 7PET, are characterized by two specific MGEs, known as *Vibrio* seventh pandemic islands I and II (VSP-I, VSP-II) [19, 20, 29, 30]. These genetic markers are pivotal for differentiating these strains from those responsible for previous pandemics.

Although the mobilome of pandemic *V. cholerae* strains has been extensively characterized, the genetic diversity within non-pandemic isolates, whether pathogenic or environmental, is still not well-understood. Previous studies primarily used PCR-based methods, focusing on confirmatory-rather than discovery-driven analyses [31–35]. In contrast, only a few studies have utilized whole-genome sequences (WGS) for a more comprehensive investigation of individual genomic islands in environmental *V. cholerae* isolates [36, 37].

In this study, we explored the mobilome diversity of non-pandemic strains and compared it to that of the 7^th^ pandemic clade of *V. cholerae*. We analyzed high-quality genome sequencing data using various methods to identify genomic islands within these strains. Additionally, we employed tools to detect defense systems, which are often found in MGEs [38–42]. Our findings revealed significant variability in the horizontal gene pool of these strains, underscoring their dynamic genetic composition. Our analysis also showed that gene operons were present in regions of genomic plasticity, often in varied combinations. Notably, some of these operons are also present in genomic islands associated with pandemic strains, suggesting a potential reservoir for novel features within non-pandemic strains of *V. cholerae*.

## MATERIAL AND METHODS

### Bacterial strains and whole-genome sequences

High-quality, well-assembled whole-genome sequences of non-pandemic environmental or clinical *V. cholerae* strains were identified from existing literature. Details of the strains used are listed in Tables S1 and S2. Unannotated *V. cholerae* genomes were annotated using the prokaryotic genome annotation pipeline (PGAP, 2022-10-03. build6384) [43].

The *de novo*-assembled genome of strain MAK757 was sequenced using PacBio technology, as described [44]. The sequencing data have been deposited in NCBI under BioProject accession number PRJNA1028811. The whole genome sequences are available in GenBank under accession numbers CP159790 and CP159791 for chromosome 1 and chromosome 2, respectively. The raw reads are available from the Sequence Read Archive (SRA) under submission numbers SRS21856092.

### Pangenome and phylogenetic reconstructions

The pangenome of all *V. cholerae* strains and *V. mimicus* was reconstructed using the PPanGGOLiN (v. 1.2.74) pipeline (all options, default parameters, sequences and annotation provided) [45]. Identified core genes were used for the subsequent phylogenetic reconstruction. To find the optimal model for each of the 1,532 core genes, ModelFinder [46] was used. The phylogeny of the 45 *V. cholerae* strains, with *V. mimicus* as an outgroup, was reconstructed using iqtree2 software (v2.2.0) [47]. The model for each individual gene was implemented from ModelFinder, and 100 bootstraps were computed in iqtree2.

### Detection of mobile genetic elements and antiviral defense systems

#### Genomic island identification

To detect MGEs, general patterns of genomic plasticity were analyzed using PanGGoLiN [45] based on the 45 genomes included in this study. Regions of genomic plasticity (RGPs) (panRGP function, with default options) [48] were thereby identified based on comparative genomics. Individual RGPs (*i.e.*, genomic islands identified in each genome), were then validated using dedicated tools that predict genomic islands based on their sequence composition (see Table S3). All programs were run with default parameters. For each genome, five different genomic island searches were performed with different assumptions underlying each individual search (for more details see [49]). The results were subsequently visualized in Geneious Prime (v. 2022, 2023 and 2024; https://www.geneious.com) and manually curated. To be considered as a *bona fide* genomic island in the context of this study, the island had to fulfill several criteria: (i) to be detected by panRGP and (ii) confirmed by at least two among the other software tools used in this study (see Table S3); and (iii) to be at least 10 kb in length, as previously suggested [6, 12]. To have a broader overview of the defensome [50] diversity, an exception to this rule was made for islands encoding predicted antiviral defense systems.

PGAP annotations and BLASTp searches were used to characterize gene clusters found in each genomic island.

#### Defense systems identification

To identify antiviral defense systems in the genomes that were included in our dataset, we used the tools DefenseFinder (accessed in April 2024) [51, 52] and PADLOC (v2.0.0) [53–55]. Identified defense systems were visualized using Geneious Prime (v. 2024; https://www.geneious.com).

#### Prophage identification

The online web tool PHASTER [56] was used to identify putative prophage sequences within the genomes.

### Alignment of genomic islands

For each insertion site in which genomic islands were detected, nucleotide sequences of the islands and neighboring genes were extracted. The comparison of the islands’ gene content and organization was visualized using clinker (v 0.0.25, default options) [57]. Nucleotide sequences of islands from pandemic strains were extracted from the genomes and aligned using MAFFT (auto option, v.7508) [58]. Each alignment was subsequently imported into Geneious Prime (v. 2022 and 2023; https://www.geneious.com) and distance matrices were calculated. Heatmaps of these matrices were generated in GraphPad Prism version 8.0.0 for MacOS (GraphPad Software, USA, www.graphpad.com).

## RESULTS & DISCUSSION

### Strain selection and phylogenetic analysis

For our analysis, we compiled a total of 46 whole-genome sequences, consisting of 45 *V. cholerae* strains and one *V. mimicus* strain, which served as an outgroup for rooting the phylogenetic tree (Tables S1 and S2). This collection encompassed genomes from eight 7PET strains (Table S1), which were used as controls. These 7PET strains were isolated at various stages of the ongoing 7^th^ cholera pandemic: early strains from the 1970s in Bangladesh and Bahrain (N16961, P27459, and E7946) [44, 59–62]; two strains from the 1990s Peruvian cholera outbreak (A1552 and C6706) [44, 60, 63, 64]; and three recent strains from 2009-2012 isolated in the Democratic Republic of the Congo (DRC193A, DRC052, and DRC072) [65]. Additionally, the dataset included a pre-pandemic strain, MAK757, isolated in 1937 in Indonesia [66], considered to be part of the so-called El Tor progenitor group [18, 67]. The assembled whole-genome sequence of this strain was deposited as part of this work (see Material and Method for details).

Our phylogenetic analysis successfully distinguished the 7PET clade as a strongly supported monophyletic group (Fig. 1a). As expected, the pre-pandemic MAK757 strain was positioned as progenitor of the El Tor group [18, 67]. In contrast to the distinct lineage of the pandemic clade, non-pandemic strains did not form a monophyletic clade. Furthermore, the analysis did not reveal any specific branching pattern based on source of isolation, indicating that strains from environmental or clinical sources did not cluster distinctly. Also, no clear biogeographic signal was observed in the non-pandemic strains’ groupings. However, a highly supported clade comprising strains originating from California (namely SA7G, E7G, SA10G, SP6G, DL4211, W7G, W6G, TP, SP7G, L6G, and SL6Y) was noted, although this did not form a monophyletic grouping with other California isolates such as SL5Y, SA5Y, and SIO. This phylogenetic resolution aligns with previous studies on the California strains that used comparative genome hybridization on microarrays [68, 69], confirming the well-supported clades A, B, and D, and revealing the polyphyletic nature of clade C.

**Figure 1.**
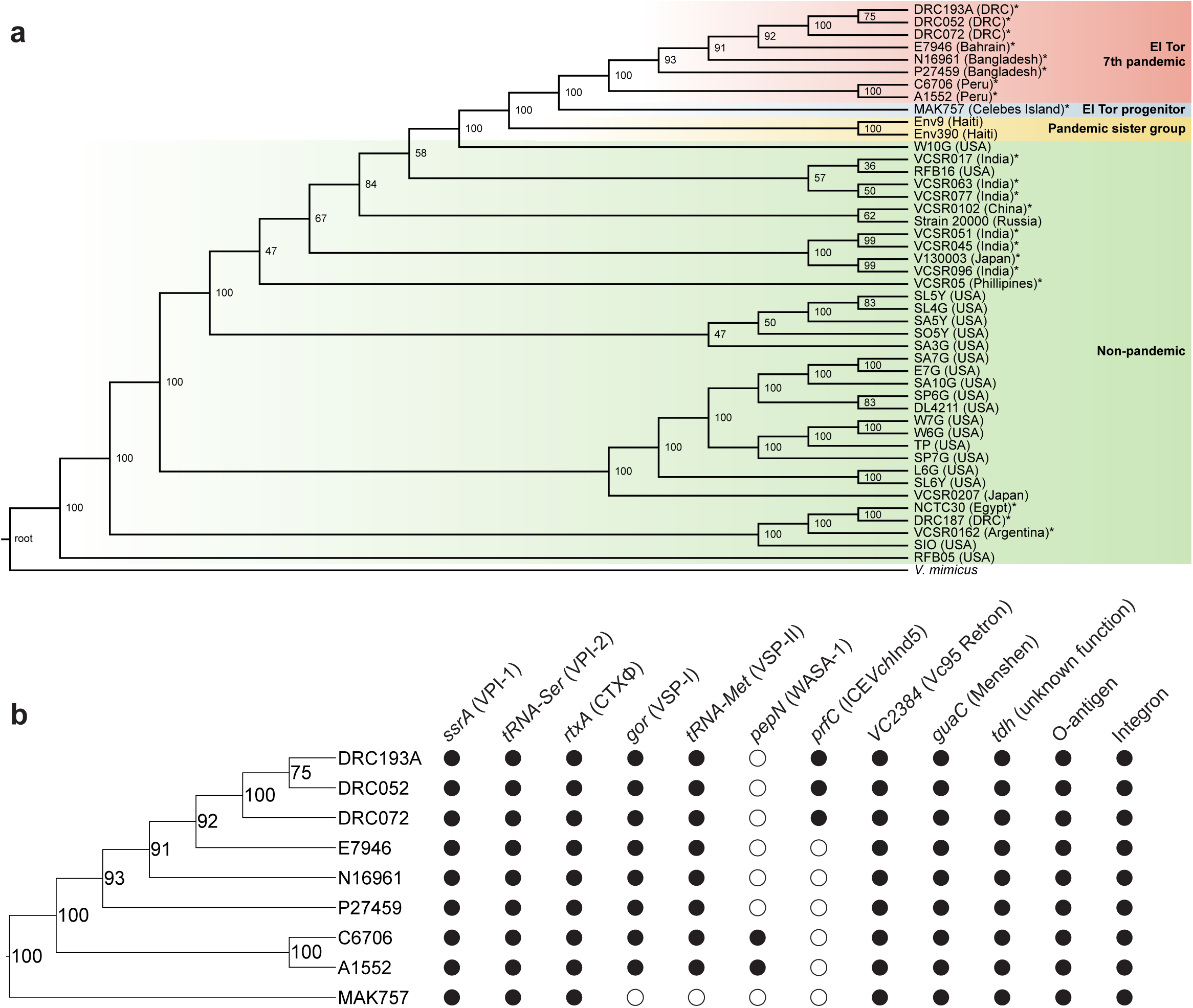
Identification of MGEs in pandemic *V. cholerae* strains. **a)** This phylogenetic tree, based on 1,532 core genes, depicts the evolutionary relationship among 45 *V. cholerae* strains, using *V. mimicus* (ATCC33655) as an outgroup. Statistical confidence was assessed with 100 bootstraps. 7PET strains (highlighted in red) and the pre-pandemic strain MAK757 (in blue), are clearly marked, whereas non-pandemic strains, including a pandemic sister group, are indicated in green and yellow, respectively. Strains isolated from patients are marked with an asterisk. **b)** A focused view of the phylogenetic tree showing only the pandemic strains. Black circles indicate the presence, and white circles the absence, of known pandemic genomic islands in each strain. The insertion sites are indicated at the top, with the names of the mobile genetic elements displayed in brackets.

Intriguingly, strain RFB05, initially identified in NCBI as a *V. cholerae* strain (Genbank: GCA_008369625.1), demonstrated phylogenetic affinity towards the root and *V. mimicus*. A secondary phylogenetic analysis (genomes used are depicted in Table S4) suggested that strain RFB05 is likely misidentified and belongs to the *V. tarriae* [70] species rather than to *V. cholerae* (Fig. S1a).

### Determination of conserved islands in pandemic strains as proof of concept

We next sought to determine the strains’ genomic islands, as outlined in the methods section. As a demonstration of our methodology’s effectiveness, we were able to identify all well-known MGEs associated with pandemic *V. cholerae* strains, as shown in Figure 1b. Notably, the pre-pandemic strain MAK757 lacked VSP-I and II, aligning with expectations (Fig. 1b). Sequence alignments and gene content analyses confirmed the expected high level of conservation of these genomic islands (*i.e.* VPI-1 and 2, VSP-I and II) across strains from various stages of the 7^th^ cholera pandemic (Fig. S1 and S2). Interestingly, DRC072 and DRC052 strains carry an additional VSP-I copy on chromosome 2, near the *uhpC* gene (encoding a MFS transporter family glucose-6-phosphate receptor; Fig. S1e).

Our methods also detected MGEs specific to certain pandemic strains, like the WASA-1 prophage in South American strains (A1552 and C6706) integrated into the alanine aminopeptidase gene (*pepN*) [18, 71] (Fig. 1b and Fig. S2e), and STX-like integrative conjugative element (ICE*Vch*Ind5; GQ463142) in DRC strains (Fig. 1b and Fig. S2f). SXT-like ICEs are common in recent clinical isolates from Asia and Africa [72] and integrate site-specifically into the *prfC* gene [73].

In our study, we identified several additional MGEs that showed a high degree of conservation across all pandemic *V. cholerae* strains (Fig. 1b). Notably, the Vc95 Retron, inserted next to gene *VC2384*, was almost identical in all examined pandemic strains (Fig. S2g). This retron plays a role in antiviral defense, comprising genes for a reverse transcriptase (RT), an ATPase, and an HNH endonuclease in addition to the non-coding RNA (ncRNA) *msr-msd* [74, 75]. Moreover, every pandemic strain was found to harbor a compact island equipped with genes encoding a predicted Menshen defense system near the *guaC* gene (Fig. S2h), and another island populated with numerous hypothetical genes near the *tdh* gene (Fig. S2i), both of them located on chromosome 2. The specific insertion sites, *VC2384*, *guaC* and *tdh,* also contain MGEs in non-pandemic strains, as discussed below.

### Mobilome diversity in non-pandemic *V. cholerae*

Shifting focus to non-pandemic *V. cholerae*, our analysis revealed the presence of MGEs across 32 insertion sites on chromosome 1 and 14 sites on chromosome 2 (Fig. 2a-b; Tables S5 and S6). Interestingly, all insertion sites housing MGEs in pandemic strains were also occupied in non-pandemic strains, with the exception of *pepN* (carrying the WASA-1 prophage in South American pandemic strains) (Fig. 2a). The most frequently occupied insertion sites in non-pandemic strains included the genes *rjg*, *ssrA*, *VC2384,* and *tRNA-Ser* in chromosome 1, as well as the integron, *tdh*, and *guaC* in chromosome 2 (Fig. 2c-d; Table S7). Specifically, while our study did not focus on these genetic elements, our analysis detected the O-antigen cluster located near the *rjg* gene on chromosome 1 [76] and a sizable integron on chromosome 2 (Fig. 2c-d). Additionally, the tmRNA gene *ssrA*, a common target for MGE integration due to independent evolution of many integrases towards this site [77], was found to be occupied by MGEs in all examined non-pandemic strains. Similarly, tRNA genes, favored by many MGEs for their conserved sequences [78], saw the *tRNA-Ser* site occupied by MGEs in over half of our non-pandemic dataset (Fig. 2c).

**Figure 2.**
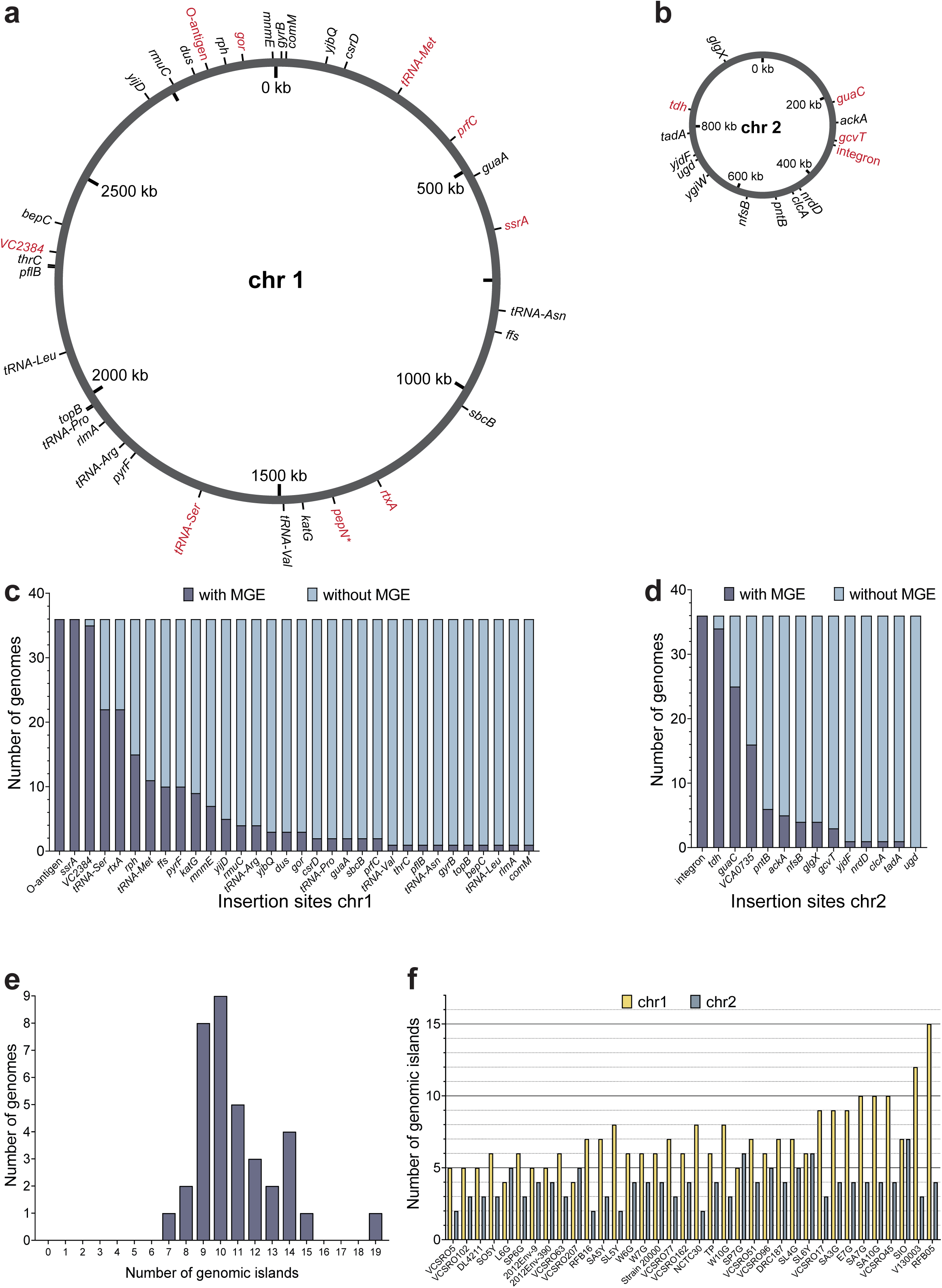
Regions of genomic plasticity in non-pandemic *V. cholerae* strains. a-b) Scheme of *V. cholerae* chromosomes 1 **(a)** and 2 **(b)**, displaying all insertion sites where MGEs are found in at least one strain. Insertion sites occupied in pandemic strains are marked in red, with those unique to pandemic strains indicated by an asterisk. The layout of these sites follows the genomic structure of pandemic strain A1552. For comprehensive details on all insertion sites, see Tables S5 and S6. **c-d)** Charts showing the presence of MGEs at insertion sites across chromosome 1 **(c)** and chromosome 2 **(d)** within the dataset of non-pandemic *V. cholerae* strains. For each site, the charts display the count of genomes with or without an MGE. **e)** The distribution graph showing the total count of genomic islands identified per genome in our non-pandemic strain dataset, highlighting *V. tarriae* RFB05 with 19 genomic islands as an outlier. **f)** Graph illustrating the quantity of MGEs located on chromosomes 1 and 2 for each non-pandemic *V. cholerae* strain examined.

MGE abundance varied from 7 to 15 genomic islands per genome, with most strains containing 10 genomic islands (taking into account the specific cutoffs we set out to consider a *bona fide* genomic island) (Fig. 2e). Most of the islands were found on the larger chromosome 1 (3 Mbp compared to 1 Mbp of chromosome 2), but some strains (L6G, VCSRO207, SP7G) had more genomic islands on chromosome 2 (Fig. 2f). This distribution pattern underscores the complex nature of genomic plasticity within *V. cholerae*, revealing the potential for diverse genetic configurations across different strains.

### Secretion systems encoded in *V. cholerae*’s mobilome

Our findings highlight a remarkable variability in gene content among MGEs located across a wide range of insertion sites within *V. cholerae* genomes. These MGEs harbor gene clusters involved in a spectrum of biological processes, from host-pathogen interactions and defense mechanisms to strategies for interbacterial competition, metabolism, and the production of cellular appendages (Fig. S3-S12).

Such diversity is indicative of the crucial role these gene clusters play in the environmental adaptation of *V. cholerae*, enabling the bacterium to navigate the myriad of selective pressures encountered in its habitats. Interestingly, the presence of these gene clusters in non-pandemic strains, some of which have been isolated from human patients, suggests their potential significance in mediating interactions with the human host. This observation underscores the complex ecology of *V. cholerae*, which spans both environmental reservoirs and human populations.

A notable finding was the widespread presence of secretion systems encoded within these genomic islands. Secretion systems, ranging from Type I to Type XI (T1SS to T11SS), are frequently found in elements that can be horizontally transferred between organisms [8]. These systems are composed of multiprotein complexes that transport a variety of substrates, including proteins and DNA, out of the cell. Such substrates are crucial for bacterial interaction with their surroundings, playing key roles in environmental adaptation and response [79]. Our data revealed a significant number of strains equipped with auxiliary T6SS and/or a T3SS gene clusters.

The T6SS functions as a sophisticated antibacterial and anti-eukaryotic arsenal, with its architecture in *V. cholerae* encoded by at least three gene clusters [80]. The primary large cluster encodes the structural framework of the system, whereas the two auxiliary clusters, Aux1 and Aux2, equip the system with Hcp and VgrG proteins that form the spear-like apparatus, complemented by adaptor proteins and effector/immunity (E/I) pairs for precise target engagement and protection against self-intoxication [81]. Pandemic *V. cholerae* strains possess an additional auxiliary cluster, Aux3, enhancing their competitive edge with an extra set of E/I genes [82–84]. Our analysis effectively pinpointed the presence of the Aux3 cluster across all examined pandemic strains, located at the *gcvT* insertion site, notable for its relatively small size under 10 kb. Their shorter size would exclude these islands from our analysis, but we kept them nonetheless for comparative reasons due to the identification of the larger (37-kb in length) version of this island, exhibiting mobile and prophage-like characteristics in non-pandemic strains 2012Env-9, 2012Env-390 and Strain 20000 (Fig. 3 and Fig. S3). This finding aligns with a previous report on Aux3 [84]. Moreover, a significant portion of our dataset included strains containing the T6SS Aux4 and 5 clusters [36, 85], reinforcing previous work [86]. Aux4 clusters were found in islands inserted at *ssrA* (Fig. S4) and *tRNA-Ser* (Fig. 4a). Aux5 clusters, on the other hand, were found in MGEs inserted near the *tdh* or *pntB* genes on chromosome 2 (Fig. S5 and S6).

**Figure 3.**
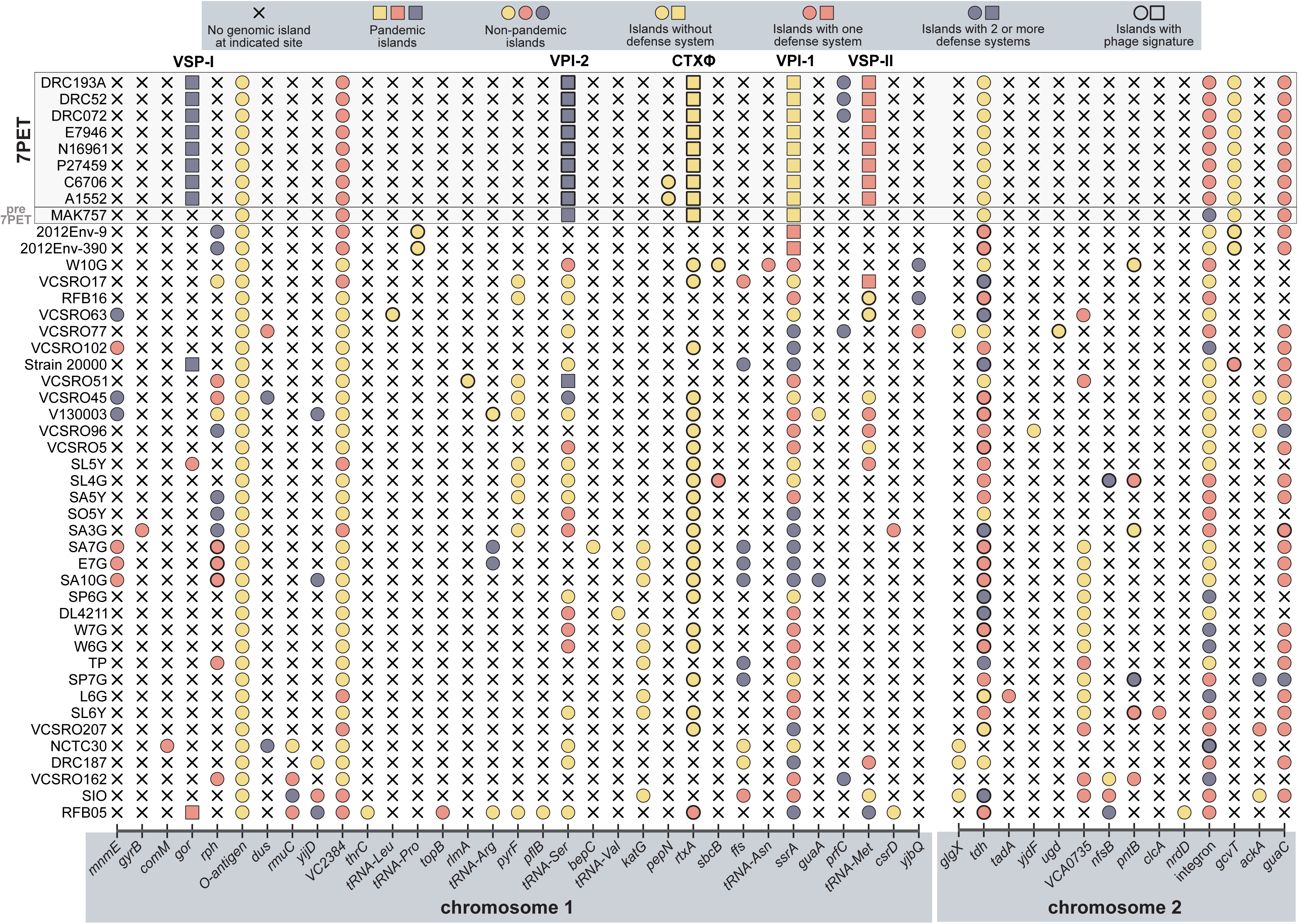
Distribution of genomic islands and antiviral defense mechanisms in *V. cholerae*. This figure maps out the locations of MGEs across *V. cholerae* strains, both pandemic and non-pandemic. The x-axis indicates the specific insertion sites where MGEs have been identified. Strain alignment on the left follows the order presented in the phylogenetic tree from Figure 1. Each circle represents a genomic island, with pandemic-specific islands marked by squares. Color-coding identifies islands without antiviral defense systems (yellow) and those with one or more defense systems (in red and gray). MGEs with phage-related signatures are highlighted with a bold outline.

**Figure 4.**
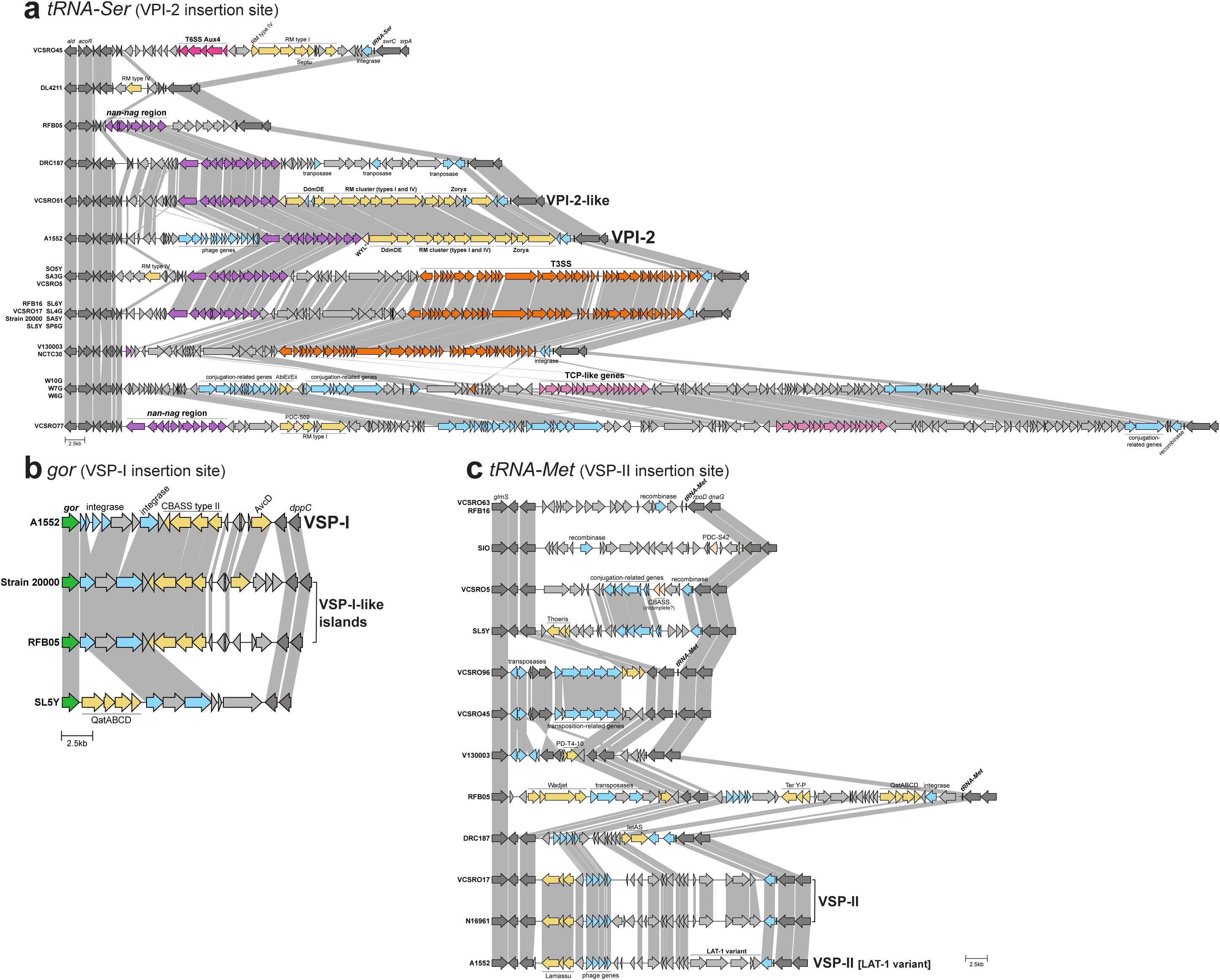
Circulation of pandemic-like islands and gene clusters in non-pandemic *V. cholerae* MGEs. Comparative gene content within genomic islands located at the *tRNA-Ser* **(a)**, *gor* **(b)**, and *tRNA-Met* **(c)** loci across various non-pandemic *V. cholerae* strains. Strains featuring these genomic islands are listed on the left. Light grey connections indicate protein identities above 30% (default parameter of the clinker pipeline). These connections are not present in the case of tRNA genes such as *tRNA-Ser* **(a)** and *tRNA-Met* **(c)**, as comparisons in clinker are based on the encoded protein sequences. Insertion sites are marked in green (arrows in the case of protein-coding genes, small squares in the case of tRNA genes). Dark grey arrows point to genes flanking the islands, with important genes labelled. Light grey arrows denote genes encoding hypothetical proteins or those not directly related to the main findings. Light blue arrows highlight mobilization-related genes. Yellow arrows mark antiviral defense genes or gene clusters, while salmon-colored arrows indicate phage defence candidates (PDC), each labelled accordingly. The gene marked with “WYL*” was identified by a loose model that detects various defense-modification systems (“Dms_other”) by PADLOC. The figure in **(a)** depicts the *nan-nag* region from VPI-2 (in purple), T3SS gene clusters (in orange) and TCP-related (VPI-1-like) clusters (in pink) distributed in non-pandemic genomic islands. T6SS Aux4 cluster is indicated in dark pink. Each panel displays the pandemic island VPI-2 **(a)**, VSP-I **(b)**, and VSP-II **(c)** of the 7PET strain A1552 for comparison.

While the T6SS is often used for interbacterial warfare, the T3SS forms a conduit between the bacterial cell and its host, facilitating the transfer of effector proteins that can alter the host’s cellular responses (for review, see [87–89]). We found that roughly one-third (13 out of the total number) of strains in our collection of non-pandemic *V. cholerae* harbored a T3SS within a genomic island located at the *tRNA-Ser* locus, a site typically associated with VPI-2 in pandemic strains. Interestingly, 11 of these 13 strains also incorporated the *nan-nag* region, a key component of VPI-2, within the same genomic island (Fig. 4a), confirming previous work [25, 90]. These findings underscore the significant role of T3SS islands in the genetic landscape of *V. cholerae*, with a notable prevalence in our dataset where over half of the strains with an island at the *tRNA-Ser* site possessed a T3SS. The distribution of these islands seems extensive, with strains isolated from various global locations over different time periods and from a range of sources (Table S2). Remarkably, one such strain, NCTC 30, was isolated from a patient in Egypt in 1916 [91], marking it as the oldest strain in our dataset and illustrating the long-standing presence of T3SS within the *V. cholerae* species.

### Circulation of pandemic-like islands in non-pandemic strains

Our analysis revealed the presence of nearly identical VPI-1 islands in non-pandemic strains 2012Env-9 and 2012Env-390, isolated from Haiti in 2012 (Fig. 3 and Fig. S4; Table S2) [92, 93]. The islands in these non-toxigenic O1 environmental isolates, previously considered as part of a pandemic sister group (Fig. 1a) [67], showed remarkable similarity to the *bona fide* VPI-1 found in pandemic strains but additionally carried genes encoding a Dnd antiviral defense system alongside the TCP and Acf clusters (Fig. S4). Interestingly, the Dnd gene cluster, but not the TCP/Acf clusters, was frequently identified within genomic islands located at the *ssrA* site in other environmental *V. cholerae* strains (L6G, TP, W10G, W7G, W6G; Fig. S4).

A similar instance was observed for a VPI-2-like island in strain VCSRO51, an O51 serogroup strain from India (isolated in 1973) [76]. This strain lacks the phage-related genes found in the *bona fide* VPI-2 (Fig. 3 and 4a). Additionally, the VPI-2-like island in VCSRO51 shows significant disruptions in genes associated with defense systems. Notably, the *ddmD* gene, a component of the DdmDE DNA defense module [26], is disrupted by two short transposase-related genes. Furthermore, within the predicted Zorya antiphage system gene cluster, *zorABCD*, there are gene insertions that encode for transposase-related proteins or proteins of unknown function (Fig. 4a). We also noticed a distribution of the sialic acid metabolism-related *nan-nag* region, an integral part of VPI-2 [24], in other non-pandemic strains, pointing to the spread of critical pandemic genetic components across different genomic backgrounds (Fig. 4a).

Further analysis identified non-pandemic strains with VSP-I-like islands, such as strain 20000, an O1 environmental isolate from Russia (2016), and the *V. tarriae* strain RFB05 from a lake in the USA (2017), both showcasing slight variations in their genomic makeup compared to VSP-I of pandemic strains (Figs. 3 and 4b), with the latter strain lacking the antiviral cytidine deaminase gene *avcD* [94]. Lastly, strain SL5Y, isolated from the Californian coast in 2004 [68], contains an island at the *gor* locus, similar to VSP-I and characterized by shared integrases. However, the gene content of this island is unique compared to other strains, notably including a gene cluster encoding a predicted QatABCD antiviral defense system among other distinctive features (Fig. 4b). These are the only occurrences of islands inserted at the *gor* site in our dataset.

We identified a VSP-II genomic island in strain VCSRO17, an O17 serogroup isolate from a 1968 patient in India, that is 99.7% identical to VSP-II from pandemic strains of the 1970s, such as N16961 (Figs. 3 and 4c). This observation underscores that mobile genetic elements can be shared between pandemic and non-pandemic strains of *V. cholerae*. Furthermore, in ten other non-pandemic strains, including *V. tarriae* RFB05, we identified islands located next to the *tRNA-Met* gene, the site in which VSP-II is integrated in 7PET strains (note that the annotation for this tRNA gene was retained as *tRNA-Met*, as initially proposed for the reference genome of strain N16196 [59], despite predictions by the PGAP [43] and tRNAscan-SE [95] software tools identifying the locus as *tRNA-Ile*). Some of these islands contain gene blocks similar to those found in pandemic VSP-II, including integrase and specific phage-like and hypothetical genes, as observed in strains DRC187 and RFB05 (Fig. 4c). However, the overall composition of these islands shows significant diversity.

Furthermore, islands found to be well-conserved among pandemic strains, though not recognized as “pandemic islands”, were detected in non-pandemic strains. For instance, the Vc95 Retron found inserted next to *VC2384* in pandemic strains (Fig. 1b and Fig. S2g) was also detected in strains 2012Env-9 and 2012Env-390 (Fig. 3 and Fig. S7). Moreover, the exact island found next to *tdh* in chromosome 2 of pandemic strains (composed of hypothetical genes) was also found in non-pandemic strains W10G and VCSRO51, and a variation of this island was encountered in strain SO5Y (Fig. S5).

Remarkably, we did not find primary pandemic islands located outside their known positions (*ssrA*, *tRNA-Ser*, *gor*, and *tRNA-Met*), with the exception of an extra VSP-I copy found on chromosome 2 in two strains from the DRC, as previously noted (Fig. S1e). However, within our dataset, we identified four strains that possess TCP-like gene clusters in islands adjacent to *tRNA-Ser*, a location different from *ssrA* where the VPI-1 (or TCP) island is typically found in pandemic strains (Fig. 4a). Specifically, strain VCSRO77, isolated from a patient in India in 1976 [76], harbors a 135 kb-long island near *tRNA-Ser* that includes both a TCP-like cluster (as in VPI-1) and a *nan-nag* region (as in VPI-2) (Fig. 4a). This type of data was not captured in earlier PCR-based genetic screenings of *V. cholerae* samples collected worldwide, which had identified environmental strains carrying key genes like *tcpA* (encoding the major TCP pilin) and/or *toxT* (encoding the virulence regulator ToxT) [31–33]. This finding within our relatively limited dataset illustrates a scenario where pandemic-like gene clusters coexist within the same genomic island, underscoring (i) the potential role of non-pandemic strains as repositories for both virulence-associated and unrelated gene clusters, and (ii) the potential emergence of hazardous combinations of disease-related gene clusters within MGEs.

### Antiviral defense systems in MGEs

Bacteriophages are notably prevalent in marine environments, where they are estimated to cause 20-40% of bacterial mortality [96]. The significant impact of phages on bacterial populations has been increasingly recognized, thanks to advances in computational biology and genome analysis [97]. These developments have revealed the broad distribution of phages and highlighted the variety of antiviral defense mechanisms that bacteria possess [98]. Contrary to earlier beliefs that such defenses were limited mainly to CRISPR-Cas and restriction-modification (RM) systems, recent research has dramatically broadened this view [40, 98–105]. It has revealed a vast array of antiphage defense strategies within bacterial genomes, now including over a hundred distinct mechanisms. This broadening of known antiviral systems reflects the dynamic evolutionary arms race between bacteriophages and their bacterial hosts.

Antiviral systems in bacteria are primarily found within MGEs [38–42, 99, 106–108]. This strategic positioning allows for the rapid exchange of defense mechanisms, essential in the bacterial-phage evolutionary arms race [101, 105, 106, 108]. In our exploration of non-pandemic *V. cholerae*, we delved into the diversity and abundance of antiviral defense mechanisms - collectively referred to as the “defensome” [50].

### Defensome diversity

In our analysis of non-pandemic *V. cholerae* strains, we quantified the prevalence of various antiviral defense systems, as depicted in Figure 5a. Consistent with previous research [51], RM systems emerged as the most prevalent defense mechanism, with 30 of the genomes (representing 83.3% of our dataset) containing at least one of these systems. Following closely, CRISPR-Cas systems were identified in 16 genomes, accounting for 47.2% of the strains we studied, despite their consistent absence from 7PET strains [109]. CRISPR-Cas Type I-F were particularly prevalent, found in islands inserted in *ssrA* and *ffs*, for example (Figs. S4 and S8). This prevalence aligns with the observations of McDonald and colleagues, who reported that Type I-F is the dominant CRISPR-Cas system within the Vibrionaceae family [110], indicating a significant trend across a broad spectrum of species.

**Figure 5.**
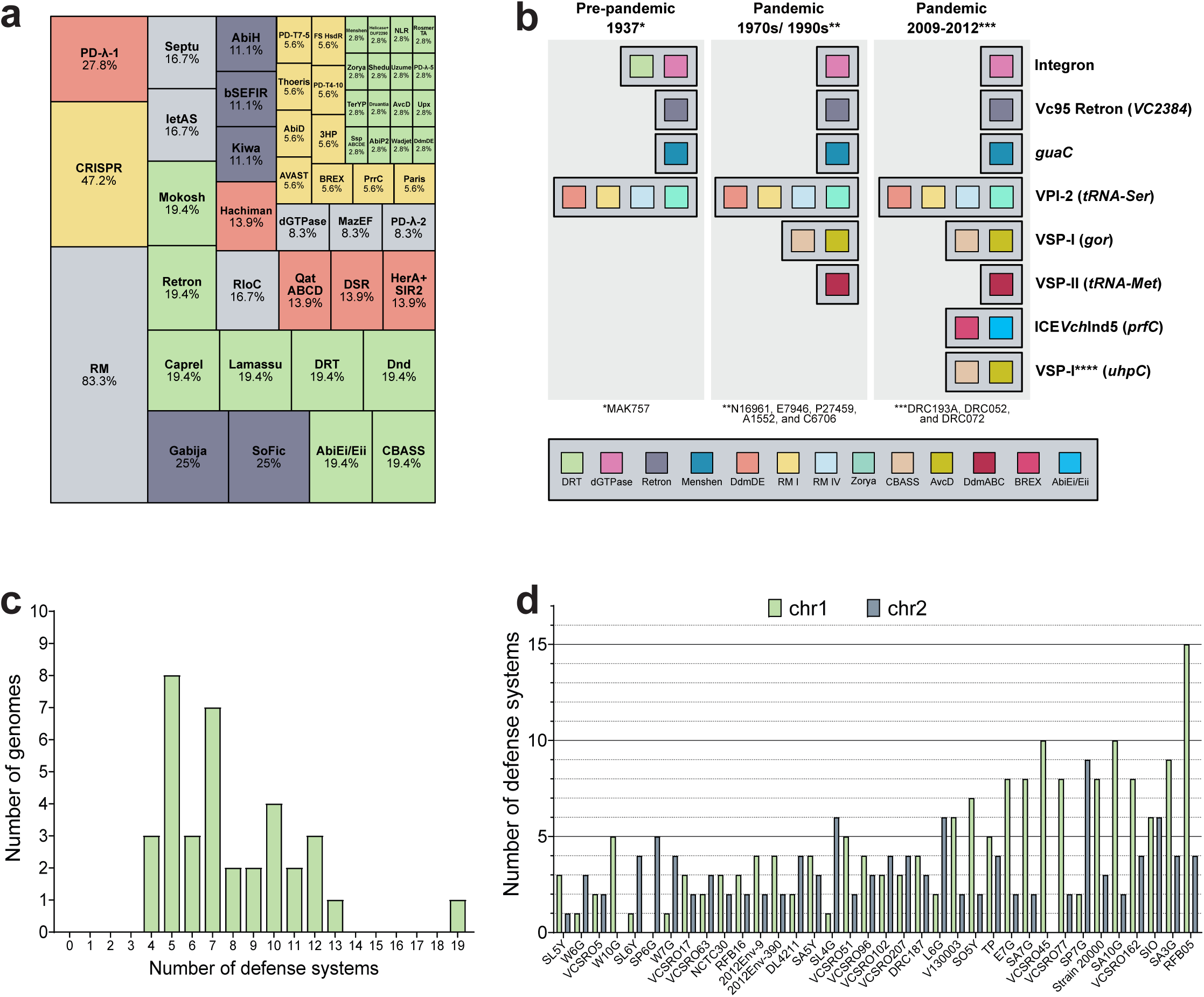
Diversity and richness of antiviral defense systems in *V. cholerae.* **a)**Frequency of antiviral defense systems identified across genomes of non-pandemic *V. cholerae* in the current dataset. **b)** This figure represents defense systems found in the pre-pandemic strain MAK757 (left), pandemic strains isolated in the 1970s/1990s (center) and in 2009-2012 (right). Each square represents a defense system, following the color code depicted in the legend. Squares are included in rectangles representing the MGE in which they are found, which are indicated on the right of the figure. **** the second VSP-I copy inserted next to *uhpC* is only found in strains DRC052 and DRC072. **c)** Overview of the number of antiphage defense systems across non-pandemic *V. cholerae* strains, including the notable case of *V. tarriae* RFB05. **d)** Detailed comparison of defense system counts on chromosomes 1 and 2 across individual non-pandemic *V. cholerae* isolates.

Other notable defense mechanisms identified include PD-λ-1, found in 10 genomes (27.7%); while Gabija and SoFic were detected in 9 genomes (25% of our dataset). Several defense systems were detected in nearly one fifth of the strains (7 genomes or 19.4%): AbiEi/Eii, CBASS, CapRel, Lamassu, DRT, Dnd, Retron and Mokosh. IetAS, Septu, and RloC were identified in 6 genomes (16.6%). Additionally, we detected 36 distinct defense systems scattered across the genomes, though each was present in five or less strains (Fig. 5a). Importantly, using the newest version of PADLOC we detected a myriad of “phage defence candidates” (PDCs) and “Hma (for Helicase-Methylase-ATPase system)-embedded candidates” [55]. These candidates were not considered as *bona fide* defense systems in our analyses, but their presence was incorporated in all figures throughout the manuscript. This varied distribution underscores the extensive diversity and specificity of antiviral defense mechanisms within non-pandemic *V. cholerae*, highlighting the species’ complex evolutionary adaptations to threats by phages and MGEs.

### Comparative analysis of defense systems in pandemic and non-pandemic V*. cholerae* strains

In pandemic *V. cholerae* strains, there is a noticeable conservation of antiviral systems across strains sharing identical genomic islands. For example, the pre-pandemic strain MAK757, dating back to 1937 and lacking VSP islands, contains seven predicted antiviral systems (Fig. 5b). In contrast, strains from the 1970s and 1990s, such as N16961, E7946, P27459, A1552, and C6706, are equipped with nine predicted systems (Fig. 5b). The most recently isolated strains from the DRC contain eleven (DRC193A) or thirteen (DRC052 and DRC072) systems (Fig. 5b). The latter group’s additional defense systems are incorporated within their ICE (ICE*Vch*Ind5) located at the *prfC* locus, as previously demonstrated for such elements of Bangladeshi *V. cholerae* isolates [39]. DRC052 and DRC072 contain an additional VSP-I encoded next to *uhpC,* as previously mentioned (Fig. S1e, Fig. 5b). This pattern highlights the evolutionary conservation of antiviral defenses in pandemic lineages and the introduction of novel defense systems through the incorporation of MGEs in the flexible genome.

In contrast, in our examination of non-pandemic *V. cholerae* strains, we discovered a broad spectrum of defense system counts per genome, ranging from 4 to 13 (with the exception of *V. tarriae* RFB05 strain, which possesses 19 systems) (Fig. 5c). The majority of strains in our study hosted approximately 5 antiviral systems each, which is in line with previous surveys reporting an average of 5 defense systems per organism based on a vast dataset of 21,738 prokaryotic genomes [51]. We also observed a significant variation in the distribution of these defense systems across chromosomes (Fig. 5d). While certain strains localized all their defense mechanisms to either chromosome 1 or 2 (as seen in strains SP6G, W10G, and VCSRO45), others exhibited a more scattered arrangement, incorporating defense systems across both chromosomes without a clear pattern. This variability suggests a complex evolutionary landscape for the acquisition and distribution of antiviral defense mechanisms in non-pandemic *V. cholerae* strains, reflecting their adaptive strategies to diverse phage challenges in their environments. Overall, *V. cholerae* showcases a wide array of defense mechanisms, positioning it among species with “many diverse systems,” similar to *Pseudomonas aeruginosa* as noted by Tesson and colleagues [51]. This diversity, even among closely related strains, underscores the importance of HGT in disseminating antiviral systems and of the “pan-immune system” concept of bacteria, as proposed by Bernheim and Sorek [101]. Indeed, our research highlights *V. cholerae*’s vast genomic diversity, particularly in its antiviral defense mechanisms. This diversity also supports the notion proposed by Hussain *et al.* in their study on *V. lentus*, where phage resistance plays a critical role in steering the diversification of the species’ mobilome [106].

### Defensome hotspots

In our subsequent analysis, we aimed to pinpoint potential genomic hotspots for the insertion of defense systems. To achieve this, we mapped the occurrence of genomic islands either possessing or lacking a defense system, across the insertion sites on chromosomes 1 and 2 (as illustrated in Figures 6a and 6b). We identified several insertion sites that were frequently occupied across multiple strains but did not consistently contain islands with known defense systems. These include the O-antigen biosynthesis cluster and the sites next to *rtxA, pyrF*, and *katG* on chromosome 1 (Fig. 6a). It is important to note that the precision of our detection of defense systems heavily relied on the capabilities of the tools we used, namely DefenseFinder and PADLOC [51–55]. This reliance suggests the possibility that additional, yet unidentified, defense systems may be encoded within these islands, awaiting discovery with more targeted analytical tools and the incorporation of new defense systems into DefenseFinder and PADLOC, as recently reported [51–55]. Indeed, as mentioned above, we identified several candidate defense systems in our dataset using the most recent version of PADLOC [55], but did not include these in our analyses due to their provisional status.

**Figure 6.**
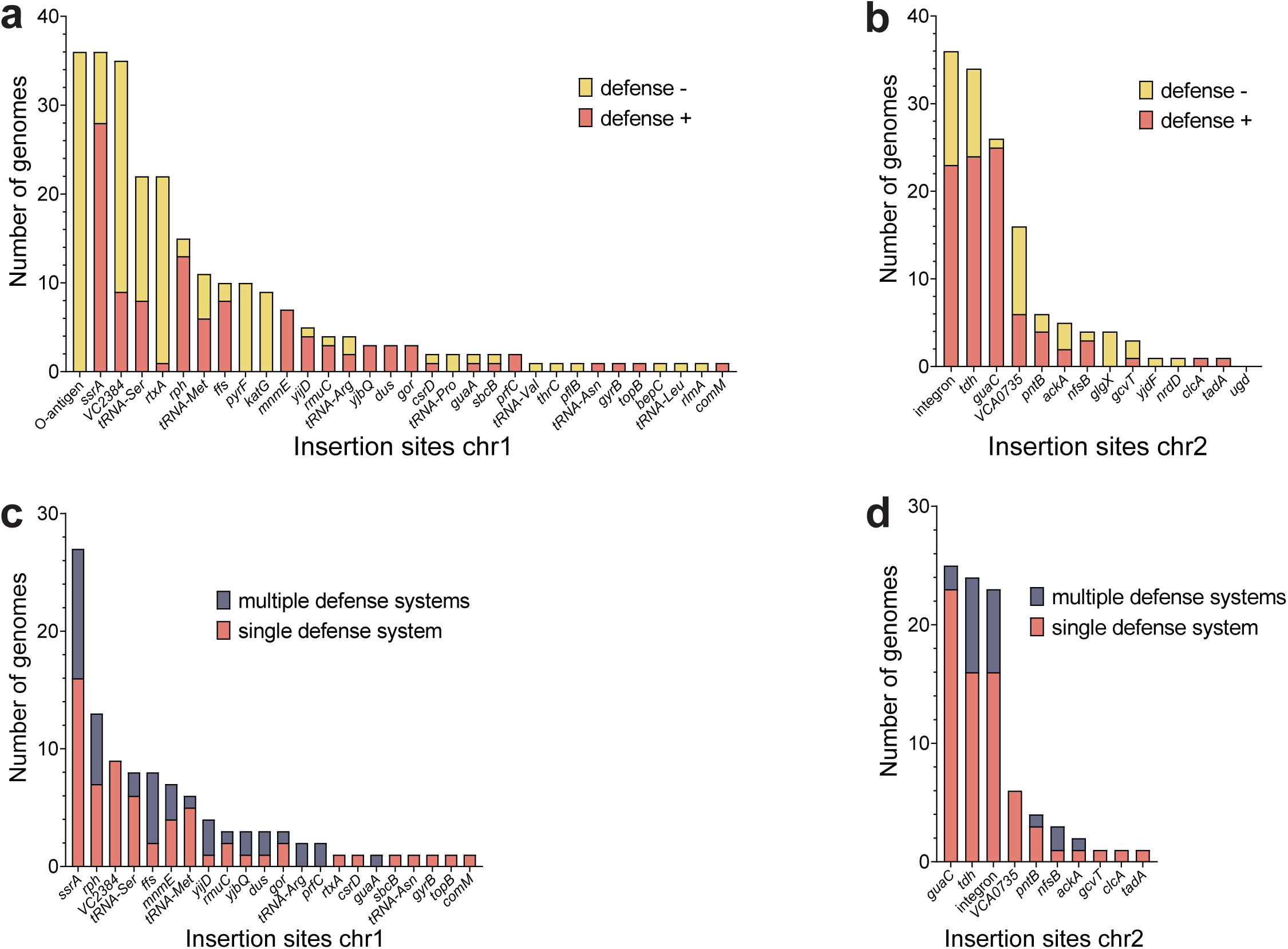
Identifying genomic regions primed for defense system insertion and the presence of defense islands in non-pandemic *V. cholerae*. a-b) Visualization of MGE occupancy on chromosomes 1 **(a)** and 2 **(b)**, distinguishing sites without defense systems (yellow) from those with one or more defense systems (red). **c-d)** Distribution of MGEs with defense systems across chromosomes 1 **(c)** and 2 **(d).** Each insertion site is plotted on the x-axis, indicating the number of MGEs containing either a single (red) or multiple (gray) defense systems.

We identified specific insertion sites that function as hotspots for antiviral defense systems, as illustrated in Figure 6. Notable examples include *ssrA* (Fig. S4), *VC2384* (Fig. S7), *ffs, yjbQ* (Fig. S8), *rph,* and *mnmE* (Fig. S9) on chromosome 1, along with *tdh* and *guaC* on chromosome 2 (Figs. S5 and S10). Among these, the *guaC* insertion site was particularly prominent, with a majority of strains harboring short MGEs equipped with defense systems (Fig. S10). Similarly, short islands inserted next to *VC2384,* where the Vc95 Retron is located in 7PET, are frequently occupied by defense systems in non-pandemic strains (Fig. S7). In many cases, however, we detected genes coding for proteins of unknown functions or those bearing domains commonly linked to defense mechanisms, including ATPases and DNA-binding proteins (Fig. S7). Several of these genes were predicted to encode phage defense candidates (PDC) by PADLOC (Fig. S7). Furthermore, our analysis revealed “hotspot regions” within genomic islands, consistently occupied by defense systems. For instance, the 3’ ends of islands next to *tdh* (Fig. S5) and near *VCA0735* (Fig. S11) on chromosome 2 showcased such patterns. Within these regions, we also detected genes similar to those found in islands next to *VC2384*, such as helicases and ATPases. This leads us to hypothesize that these genes or gene clusters might represent defense systems that current detection tools, such as PADLOC and DefenseFinder, failed to recognize, or they could be novel systems not yet characterized. A recent study on sedentary chromosomal integrons, including those in 7PET strain N16961, supports this idea [111]. Despite the absence of predicted anti-viral defense systems in these integron islands using DefenseFinder, experimental validation revealed that an astonishing 10% of integron cassettes exhibited anti-phage activity when artificially expressed in *V. cholerae* or *E. coli.* This suggests that integrons may serve as reservoirs of bacterial anti-phage defense systems [111].

### Defense islands

Defense systems in *V. cholerae* often cluster together, consistent with the concept of “defense islands” [99, 103, 106]. Our interest was piqued by two main questions: (i) the prevalence of these defense islands within our dataset, and (ii) their frequency of occurrence at specific genomic sites. To explore these aspects, we catalogued all insertion sites that hosted at least one island equipped with a defense system and analyzed the occurrence of MGEs with one or more defense systems at each site. Our findings revealed that, across most insertion sites, islands with a singular predicted defense system predominated over those with multiple systems (Fig. 6c-d), although this observation is made within the limitations of our detection capabilities, as outlined above. However, certain locations, such as *ssrA* (Fig. S4), *ffs, yjbQ* (Fig. S8), *rph,* (Fig. S9a), *yijD* and *folD* (Fig. S12a-b), on chromosome 1, stood out as exceptions, where defense islands were more commonly found.

### Prevalence of Prophage-Derived MGEs

We identified prophage signatures within our genomic dataset, encompassing both pandemic and non-pandemic strains, and determined whether these signatures coincided with the detected MGEs. We found that certain insertion sites frequently hosted MGEs bearing phage signatures. Notably, the *rtxA* site on chromosome 1, where the CTX prophage integrates in toxigenic strains, was one such location (Fig. 3). Other sites, including *tRNA-Leu, tRNA-Pro, rlmA*, *pepN*, and *sbcB* on chromosome 1, and *ugd* on chromosome 2, were exclusively associated with prophage-related MGEs, although these sites were generally not frequently occupied (Fig. 3). Additionally, we identified specific strains where MGEs of prophage origin were present at particular sites, such as *tRNA-Ser, tRNA-Met,* and *tRNA-Arg* on chromosome 1 (Fig. 4a and 4c, and S12b), and *gcvT*, *tdh, pntB,* and *guaC* on chromosome 2 (Figs. S3, S5, S6, and S10). Intriguingly, prophage signatures at the *tRNA-Ser* site were predominantly observed in VPI-2 from pandemic strains. However, ultimately, no clear correlation was found between the presence of prophage signatures and the existence of defense systems (Fig. 3).

## CONCLUSION

To our knowledge, this study represents the first comprehensive examination of the mobilome and defensome diversity within the *V. cholerae* species, extending beyond the 7PET pandemic clade. This work establishes a foundational database cataloging MGEs and antiphage defense systems identified in these strains, highlighting their distribution, frequency, and variability. Although this study includes a limited number of non-pandemic strains, our findings point toward potentially new defense systems and unexplored molecular mechanisms. These discoveries offer rich opportunities for future scientific exploration and a deeper understanding of *V. cholerae* as a species.

## ACKNOWLEDGMENTS

We thank the members of the Blokesch laboratory for fruitful discussions. We acknowledge the staff of the Lausanne Genomic Technologies Facility at the University of Lausanne and the members of the EPFL/UNIL Bioinformatics Competence Center for their contributions to sample processing, sequencing, genome assembly, and SMRT sequencing analysis. This work was supported by the Swiss National Science Foundation (310030_185022), a Consolidator Grant by the European Research Council (724630), and an International Research Scholarship by the Howard Hughes Medical Institute (HHMI) (grant 55008726) to M.B..

## AUTHOR CONTRIBUTIONS

N.C.D.D., A.L., and M.B. conceived the project and designed the details of the study; N.C.D.D. and A.L. developed the concept of the pipeline to detect genomic islands and defense systems; A.L. developed the pipeline and scripts used in this study; N.C.D.D. analyzed the data; M.B. oversaw the project’s implementation, verified findings, and secured funding; N.C.D.D, A.L., and M.B. wrote the manuscript.

## REFERENCES

1. Doolittle WF, Papke RT: Genomics and the bacterial species problem. Genome Biol 2006, 7:116.

2. Shapiro BJ, Polz MF: Ordering microbial diversity into ecologically and genetically cohesive units. Trends Microbiol 2014, 22:235–247.

3. Frost LS, Leplae R, Summers AO, Toussaint A: Mobile genetic elements: the agents of open source evolution. Nat Rev Microbiol 2005, 3:722–732.

4. Toussaint A, Chandler M: Prokaryote genome fluidity: toward a system approach of the mobilome. Methods Mol Biol 2012, 804:57–80.

5. Innamorati KA, Earl JP, Aggarwal SD, Ehrlich GD, Hiller NL: The Bacterial Guide to Designing a Diversified Gene Portfolio. In The Pangenome: Diversity, Dynamics and Evolution of Genomes. Edited by Tettelin H, Medini D. Cham (CH); 2020: 51–87

6. Arnold BJ, Huang IT, Hanage WP: Horizontal gene transfer and adaptive evolution in bacteria. Nat Rev Microbiol 2022, 20:206–218.

7. Ochman H, Lawrence JG, Groisman EA: Lateral gene transfer and the nature of bacterial innovation. Nature 2000, 405:299–304.

8. Dobrindt U, Hochhut B, Hentschel U, Hacker J: Genomic islands in pathogenic and environmental microorganisms. Nat Rev Microbiol 2004, 2:414–424.

9. Osborn AM, Böltner D: When phage, plasmids, and transposons collide: genomic islands, and conjugative- and mobilizable-transposons as a mosaic continuum. Plasmid 2002, 48:202–212.

10. Schmidt H, Hensel M: Pathogenicity islands in bacterial pathogenesis. Clin Microbiol Rev 2004, 17:14–56.

11. Ahmed N, Dobrindt U, Hacker J, Hasnain SE: Genomic fluidity and pathogenic bacteria: applications in diagnostics, epidemiology and intervention. Nat Rev Microbiol 2008, 6:387–394.

12. Juhas M, van der Meer JR, Gaillard M, Harding RM, Hood DW, Crook DW: Genomic islands: tools of bacterial horizontal gene transfer and evolution. FEMS Microbiol Rev 2009, 33:376–393.

13. Faruque SM, Albert MJ, Mekalanos JJ: Epidemiology, genetics, and ecology of toxigenic Vibrio cholerae. Microbiol Mol Biol Rev 1998, 62:1301–1314.

14. Cottingham KL, Chiavelli DA, Taylor RK: Environmental microbe and human pathogen: The ecology and microbiology of *Vibrio cholerae*. Front Ecol Environ Microbiol 2003, 1:80–86.

15. Communicable diseases following natural disasters

16. Clemens JD, van Loon F, Sack DA, Rao MR, Ahmed F, Chakrabort YJ, Kay BA, Khan MR, Yunus MD, Harris JR, et al.: Biotype as determinant of natural immunising effect of cholera. Lancet 1991, 337:883–884.

17. Robins WP, Mekalanos JJ: Genomic science in understanding cholera outbreaks and evolution of *Vibrio cholerae* as a human pathogen. Curr Top Microbiol Immunol 2014, 379:211–229.

18. Mutreja A, Kim DW, Thomson NR, Connor TR, Lee JH, Kariuki S, Croucher NJ, Choi SY, Harris SR, Lebens M, et al: Evidence for several waves of global transmission in the seventh cholera pandemic. Nature 2011, 477:462–465.

19. Boyd EF, Carpenter MR, Chowdhury N, Cohen AL, Haines-Menges BL, Kalburge SS, Kingston JJ, Lubin JB, Ongagna-Yhombi SY, Whitaker WB: Post-Genomic Analysis of Members of the Family Vibrionaceae. Microbiol Spectr 2015, 3:VE-0009-2014.

20. Dziejman M, Balon E, Boyd D, Fraser CM, Heidelberg JF, Mekalanos JJ: Comparative genomic analysis of *Vibrio cholerae*: genes that correlate with cholera endemic and pandemic disease. Proc Natl Acad Sci USA 2002, 99:1556–1561.

21. Taylor RK, Miller VL, Furlong DB, Mekalanos JJ: Use of *phoA* gene fusions to identify a pilus colonization factor coordinately regulated with cholera toxin. Proc Natl Acad Sci USA 1987, 84:2833–2837.

22. Karaolis DKR, Johnson JA, Bailey CC, Boedeker EC, Kaper JB, Reeves PR: A *Vibrio cholerae* pathogenicity island associated with epidemic and pandemic strains. Proc Natl Acad Sci USA 1998, 95:3134–3139.

23. Waldor MK, Mekalanos JJ: Lysogenic conversion by a filamentous phage encoding cholera toxin. Science 1996, 272:1910–1914.

24. Jermyn WS, Boyd EF: Molecular evolution of Vibrio pathogenicity island-2 (VPI-2): mosaic structure among Vibrio cholerae and Vibrio mimicus natural isolates. Microbiology 2005, 151:311–322.

25. Murphy RA, Boyd EF: Three pathogenicity islands of *Vibrio cholerae* can excise from the chromosome and form circular intermediates. J Bacteriol 2008, 190:636–647.

26. Jaskólska M, Adams DW, Blokesch M: Two defence systems eliminate plasmids from seventh pandemic *Vibrio cholerae*. Nature 2022, 604:323–329.

27. Vizzarro G, Lemopoulos A, Adams DW, Blokesch M: *Vibrio cholerae* pathogenicity island 2 encodes two distinct types of restriction systems. J. Bacteriol. (in press; doi: 10.1128/jb.00145-24) & bioRxiv 2024:2024.2004.2004.588119.

28. Gomez JB, Waters CM: A *Vibrio cholerae* Type IV restriction system targets glucosylated 5-hydroxyl methyl cytosine to protect against phage infection. bioRxiv 2024:2024.2004.2005.588314.

29. O’Shea YA, Reen FJ, Quirke AM, Boyd EF: Evolutionary genetic analysis of the emergence of epidemic *Vibrio cholerae* isolates on the basis of comparative nucleotide sequence analysis and multilocus virulence gene profiles. J Clin Microbiol 2004, 42:4657–4671.

30. Davies BW, Bogard RW, Young TS, Mekalanos JJ: Coordinated regulation of accessory genetic elements produces cyclic di-nucleotides for *V. cholerae* virulence. Cell 2012, 149:358–370.

31. Rivera IN, Chun J, Huq A, Sack RB, Colwell RR: Genotypes associated with virulence in environmental isolates of Vibrio cholerae. Appl Environ Microbiol 2001, 67:2421–2429.

32. Faruque SM, Kamruzzaman M, Meraj IM, Chowdhury N, Nair GB, Sack RB, Colwell RR, Sack DA: Pathogenic potential of environmental *Vibrio cholerae* strains carrying genetic variants of the toxin-coregulated pilus pathogenicity island. Infect Immun 2003, 71:1020–1025.

33. Faruque SM, Chowdhury N, Kamruzzaman M, Dziejman M, Rahman MH, Sack DA, Nair GB, Mekalanos JJ: Genetic diversity and virulence potential of environmental *Vibrio cholerae* population in a cholera-endemic area. Proc Natl Acad Sci USA 2004, 101:2123–2128.

34. Gennari M, Ghidini V, Caburlotto G, Lleo MM: Virulence genes and pathogenicity islands in environmental *Vibrio* strains nonpathogenic to humans. FEMS Microbiol Ecol 2012, 82:563–573.

35. Bernardy EE, Turnsek MA, Wilson SK, Tarr CL, Hammer BK: Diversity of Clinical and Environmental Isolates of *Vibrio cholerae* in Natural Transformation and Contact-Dependent Bacterial Killing Indicative of Type VI Secretion System Activity. Appl Environ Microbiol 2016, 18:2833–2842.

36. Labbate M, Orata FD, Petty NK, Jayatilleke ND, King WL, Kirchberger PC, Allen C, Mann G, Mutreja A, Thomson NR, et al: A genomic island in *Vibrio cholerae* with VPI-1 site-specific recombination characteristics contains CRISPR-Cas and type VI secretion modules. Sci Rep 2016, 6:36891.

37. Rapa RA, Islam A, Monahan LG, Mutreja A, Thomson N, Charles IG, Stokes HW, Labbate M: A genomic island integrated into *recA* of *Vibrio cholerae* contains a divergent recA and provides multi-pathway protection from DNA damage. Environ Microbiol 2015, 17:1090–1102.

38. Hochhauser D, Millman A, Sorek R: The defense island repertoire of the *Escherichia coli* pan-genome. PLoS Genet 2023, 19:e1010694.

39. LeGault KN, Hays SG, Angermeyer A, McKitterick AC, Johura FT, Sultana M, Ahmed T, Alam M, Seed KD: Temporal shifts in antibiotic resistance elements govern phage-pathogen conflicts. Science 2021, 373.

40. Vassallo CN, Doering CR, Littlehale ML, Teodoro GIC, Laub MT: A functional selection reveals previously undetected anti-phage defence systems in the *E. coli* pangenome. Nat Microbiol 2022, 7:1568–1579.

41. Rousset F, Depardieu F, Miele S, Dowding J, Laval AL, Lieberman E, Garry D, Rocha EPC, Bernheim A, Bikard D: Phages and their satellites encode hotspots of antiviral systems. Cell Host Microbe 2022, 30:740–753 e745.

42. Benler S, Faure G, Altae-Tran H, Shmakov S, Zheng F, Koonin E: Cargo Genes of Tn7-Like Transposons Comprise an Enormous Diversity of Defense Systems, Mobile Genetic Elements, and Antibiotic Resistance Genes. mBio 2021, 12:e0293821.

43. Tatusova T, DiCuccio M, Badretdin A, Chetvernin V, Nawrocki EP, Zaslavsky L, Lomsadze A, Pruitt KD, Borodovsky M, Ostell J: NCBI prokaryotic genome annotation pipeline. Nucleic Acids Res 2016, 44:6614–6624.

44. Stutzmann S, Blokesch M: Comparison of chitin-induced natural transformation in pandemic *Vibrio cholerae* O1 El Tor strains. Environ Microbiol 2020, 22:4149–4166.

45. Gautreau G, Bazin A, Gachet M, Planel R, Burlot L, Dubois M, Perrin A, Medigue C, Calteau A, Cruveiller S, et al: PPanGGOLiN: Depicting microbial diversity via a partitioned pangenome graph. PLoS Comput Biol 2020, 16:e1007732.

46. Kalyaanamoorthy S, Minh BQ, Wong TKF, von Haeseler A, Jermiin LS: ModelFinder: fast model selection for accurate phylogenetic estimates. Nat Methods 2017, 14:587–589.

47. Minh BQ, Schmidt HA, Chernomor O, Schrempf D, Woodhams MD, von Haeseler A, Lanfear R: IQ-TREE 2: New Models and Efficient Methods for Phylogenetic Inference in the Genomic Era. Mol Biol Evol 2020, 37:1530–1534.

48. Bazin A, Gautreau G, Medigue C, Vallenet D, Calteau A: panRGP: a pangenome-based method to predict genomic islands and explore their diversity. Bioinformatics 2020, 36:i651–i658.

49. da Silva Filho AC, Raittz RT, Guizelini D, De Pierri CR, Augusto DW, Dos Santos-Weiss ICR, Marchaukoski JN: Comparative Analysis of Genomic Island Prediction Tools. Front Genet 2018, 9:619.

50. Beavogui A, Lacroix A, Wiart N, Poulain J, Delmont TO, Paoli L, Wincker P, Oliveira PH: The defensome of complex bacterial communities. Nat Commun 2024, 15:2146.

51. Tesson F, Herve A, Mordret E, Touchon M, d’Humieres C, Cury J, Bernheim A: Systematic and quantitative view of the antiviral arsenal of prokaryotes. Nat Commun 2022, 13:2561.

52. Abby SS, Neron B, Menager H, Touchon M, Rocha EP: MacSyFinder: a program to mine genomes for molecular systems with an application to CRISPR-Cas systems. PLoS One 2014, 9:e110726.

53. Payne LJ, Todeschini TC, Wu Y, Perry BJ, Ronson CW, Fineran PC, Nobrega FL, Jackson SA: Identification and classification of antiviral defence systems in bacteria and archaea with PADLOC reveals new system types. Nucleic Acids Res 2021, 49:10868–10878.

54. Payne LJ, Meaden S, Mestre MR, Palmer C, Toro N, Fineran PC, Jackson SA: PADLOC: a web server for the identification of antiviral defence systems in microbial genomes. Nucleic Acids Res 2022, 50:W541–W550.

55. Payne LJ, Hughes TCD, Fineran PC, Jackson SA: New antiviral defences are genetically embedded within prokaryotic immune systems. bioRxiv 2024:2024.2001.2029.577857.

56. Arndt D, Grant JR, Marcu A, Sajed T, Pon A, Liang Y, Wishart DS: PHASTER: a better, faster version of the PHAST phage search tool. Nucleic Acids Res 2016, 44:W16–21.

57. Gilchrist CLM, Chooi YH: Clinker & clustermap.js: Automatic generation of gene cluster comparison figures. Bioinformatics 2021, 37:2473–2475.

58. Katoh K, Standley DM: MAFFT multiple sequence alignment software version 7: improvements in performance and usability. Mol Biol Evol 2013, 30:772–780.

59. Heidelberg JF, Eisen JA, Nelson WC, Clayton RA, Gwinn ML, Dodson RJ, Haft DH, Hickey EK, Peterson JD, Umayam L, et al: DNA sequence of both chromosomes of the cholera pathogen *Vibrio cholerae*. Nature 2000, 406:477–483.

60. Matthey N, Drebes Dörr NC, Blokesch M: Long-Read-Based Genome Sequences of Pandemic and Environmental *Vibrio cholerae* Strains. Microbiol Resour Announc 2018, 7:e01574–01518.

61. Miller VL, DiRita VJ, Mekalanos JJ: Identification of *toxS*, a regulatory gene whose product enhances ToxR-mediated activation of the cholera toxin promoter. J Bacteriol 1989, 171:1288–1293.

62. Pearson GD, Woods A, Chiang SL, Mekalanos JJ: CTX genetic element encodes a site-specific recombination system and an intestinal colonization factor. Proc Natl Acad Sci USA 1993, 90:3750–3754.

63. Yildiz FH, Schoolnik GK: Role of *rpoS* in stress survival and virulence of *Vibrio cholerae*. J Bacteriol 1998, 180:773–784.

64. Stutzmann S, Blokesch M: Circulation of a Quorum-Sensing-Impaired Variant of *Vibrio cholerae* Strain C6706 Masks Important Phenotypes. mSphere 2016, 1:e00098–16.

65. Lemopoulos A, Miwanda B, Drebes Dörr Natália C, Stutzmann S, Bompangue D, Muyembe-Tamfum J-J, Blokesch M: Genome sequences of Vibrio cholerae strains isolated in the DRC between 2009 and 2012. Microbio Resour Announc 2024, 0:e00827–00823.

66. Chattopadhyay DJ, Sarkar BL, Ansari MQ, Chakrabarti BK, Roy MK, Ghosh AN, Pal SC: New phage typing scheme for *Vibrio cholerae* O1 biotype El Tor strains. J Clin Microbiol 1993, 31:1579–1585.

67. Boucher Y: Sustained Local Diversity of *Vibrio cholerae* O1 Biotypes in a Previously Cholera-Free Country. MBio 2016, 7.

68. Keymer DP, Miller MC, Schoolnik GK, Boehm AB: Genomic and phenotypic diversity of coastal *Vibrio cholerae* strains is linked to environmental factors. Appl Environ Microbiol 2007, 73:3705–3714.

69. Miller MC, Keymer DP, Avelar A, Boehm AB, Schoolnik GK: Detection and transformation of genome segments that differ within a coastal population of *Vibrio cholerae* strains. Appl Environ Microbiol 2007, 73:3695–3704.

70. Islam MT, Liang K, Orata FD, Im MS, Alam M, Lee CC, Boucher YF: *Vibrio tarriae* sp. nov., a novel member of the Cholerae clade. Int J Syst Evol Microbiol 2022, 72.

71. Garza DR, Thompson CC, Loureiro EC, Dutilh BE, Inada DT, Junior EC, Cardoso JF, Nunes MR, de Lima CP, Silvestre RV, et al: Genome-wide study of the defective sucrose fermenter strain of *Vibrio cholerae* from the Latin American cholera epidemic. PLoS One 2012, 7:e37283.

72. Wozniak RA, Fouts DE, Spagnoletti M, Colombo MM, Ceccarelli D, Garriss G, Dery C, Burrus V, Waldor MK: Comparative ICE genomics: insights into the evolution of the SXT/R391 family of ICEs. PLoS Genet 2009, 5:e1000786.

73. Hochhut B, Waldor MK: Site-specific integration of the conjugal Vibrio cholerae SXT element into prfC. Mol Microbiol 1999, 32:99–110.

74. Millman A, Bernheim A, Stokar-Avihail A, Fedorenko T, Voichek M, Leavitt A, Oppenheimer-Shaanan Y, Sorek R: Bacterial Retrons Function In Anti-Phage Defense. Cell 2020, 183:1551–1561 e1512.

75. Bobonis J, Mitosch K, Mateus A, Karcher N, Kritikos G, Selkrig J, Zietek M, Monzon V, Pfalz B, Garcia-Santamarina S, et al: Bacterial retrons encode phage-defending tripartite toxin-antitoxin systems. Nature 2022, 609:144–150.

76. Murase K, Arakawa E, Izumiya H, Iguchi A, Takemura T, Kikuchi T, Nakagawa I, Thomson NR, Ohnishi M, Morita M: Genomic dissection of the *Vibrio cholerae* O-serogroup global reference strains: reassessing our view of diversity and plasticity between two chromosomes. Microb Genom 2022, 8.

77. Williams KP: Traffic at the tmRNA gene. J Bacteriol 2003, 185:1059–1070.

78. Williams KP: Integration sites for genetic elements in prokaryotic tRNA and tmRNA genes: sublocation preference of integrase subfamilies. Nucleic Acids Res 2002, 30:866–875.

79. Costa TRD, Felisberto-Rodrigues C, Meir A, Prevost MS, Redzej A, Trokter M, Waksman G: Secretion systems in Gram-negative bacteria: structural and mechanistic insights. Nat Rev Microbiol 2015, 13:343–359.

80. Boyer F, Fichant G, Berthod J, Vandenbrouck Y, Attree I: Dissecting the bacterial type VI secretion system by a genome wide in silico analysis: what can be learned from available microbial genomic resources? BMC Genomics 2009, 10:104.

81. Crisan CV, Hammer BK: The Vibrio cholerae type VI secretion system: toxins, regulators and consequences. Environ Microbiol 2020, 22:4112–4122.

82. Altindis E, Dong T, Catalano C, Mekalanos J: Secretome analysis of *Vibrio cholerae* type VI secretion system reveals a new effector-immunity pair. mBio 2015, 6:e00075–15.

83. Hersch SJ, Watanabe N, Stietz MS, Manera K, Kamal F, Burkinshaw B, Lam L, Pun A, Li M, Savchenko A, Dong TG: Envelope stress responses defend against type six secretion system attacks independently of immunity proteins. Nat Microbiol 2020, 5:706–714.

84. Santoriello FJ, Michel L, Unterweger D, Pukatzki S: Pandemic *Vibrio cholerae* shuts down site-specific recombination to retain an interbacterial defence mechanism. Nat Commun 2020, 11:6246.

85. Crisan CV, Chande AT, Williams K, Raghuram V, Rishishwar L, Steinbach G, Watve SS, Yunker P, Jordan IK, Hammer BK: Analysis of Vibrio cholerae genomes identifies new type VI secretion system gene clusters. Genome Biol 2019, 20:163.

86. Otto SB, Servajean R, Lemopoulos A, Bitbol AF, Blokesch M: Interactions between pili affect the outcome of bacterial competition driven by the type VI secretion system. Curr Biol 2024, 34:2403–2417 e2409.

87. Büttner D: Protein export according to schedule: architecture, assembly, and regulation of type III secretion systems from plant- and animal-pathogenic bacteria. Microbiol Mol Biol Rev 2012, 76:262–310.

88. Galan JE, Waksman G: Protein-Injection Machines in Bacteria. Cell 2018, 172:1306–1318.

89. Wagner S, Grin I, Malmsheimer S, Singh N, Torres-Vargas CE, Westerhausen S: Bacterial type III secretion systems: a complex device for the delivery of bacterial effector proteins into eukaryotic host cells. FEMS Microbiol Lett 2018, 365.

90. Carpenter MR, Kalburge SS, Borowski JD, Peters MC, Colwell RR, Boyd EF: CRISPR-Cas and Contact-Dependent Secretion Systems Present on Excisable Pathogenicity Islands with Conserved Recombination Modules. J Bacteriol 2017, 199.

91. Dorman MJ, Kane L, Domman D, Turnbull JD, Cormie C, Fazal MA, Goulding DA, Russell JE, Alexander S, Thomson NR: The history, genome and biology of NCTC 30: a non-pandemic Vibrio cholerae isolate from World War One. Proc Biol Sci 2019, 286:20182025.

92. Azarian T, Ali A, Johnson JA, Jubair M, Cella E, Ciccozzi M, Nolan DJ, Farmerie W, Rashid MH, Sinha-Ray S, et al: Non-toxigenic environmental *Vibrio cholerae* O1 strain from Haiti provides evidence of pre-pandemic cholera in Hispaniola. Sci Rep 2016, 6:36115.

93. Azarian T, Ali A, Johnson JA, Mohr D, Prosperi M, Veras NM, Jubair M, Strickland SL, Rashid MH, Alam MT, et al: Phylodynamic analysis of clinical and environmental *Vibrio cholerae* isolates from Haiti reveals diversification driven by positive selection. mBio 2014, 5.

94. Hsueh BY, Severin GB, Elg CA, Waldron EJ, Kant A, Wessel AJ, Dover JA, Rhoades CR, Ridenhour BJ, Parent KN, et al: Phage defence by deaminase-mediated depletion of deoxynucleotides in bacteria. Nat Microbiol 2022, 7:1210–1220.

95. Chan PP, Lowe TM: tRNAscan-SE: Searching for tRNA Genes in Genomic Sequences. Methods Mol Biol 2019, 1962:1–14.

96. Suttle CA: Marine viruses – major players in the global ecosystem. Nat Rev Microbiol 2007, 5:801–812.

97. Roux S, Hallam SJ, Woyke T, Sullivan MB: Viral dark matter and virus-host interactions resolved from publicly available microbial genomes. Elife 2015, 4.

98. Hampton HG, Watson BNJ, Fineran PC: The arms race between bacteria and their phage foes. Nature 2020, 577:327–336.

99. Makarova KS, Wolf YI, Snir S, Koonin EV: Defense islands in bacterial and archaeal genomes and prediction of novel defense systems. J Bacteriol 2011, 193:6039–6056.

100. Georjon H, Bernheim A: The highly diverse antiphage defence systems of bacteria. Nat Rev Microbiol 2023.

101. Bernheim A, Sorek R: The pan-immune system of bacteria: antiviral defence as a community resource. Nat Rev Microbiol 2020, 18:113–119.

102. van Houte S, Buckling A, Westra ER: Evolutionary Ecology of Prokaryotic Immune Mechanisms. Microbiol Mol Biol Rev 2016, 80:745–763.

103. Doron S, Melamed S, Ofir G, Leavitt A, Lopatina A, Keren M, Amitai G, Sorek R: Systematic discovery of antiphage defense systems in the microbial pangenome. Science 2018, 359.

104. Gao L, Altae-Tran H, Bohning F, Makarova KS, Segel M, Schmid-Burgk JL, Koob J, Wolf YI, Koonin EV, Zhang F: Diverse enzymatic activities mediate antiviral immunity in prokaryotes. Science 2020, 369:1077–1084.

105. Koonin EV, Makarova KS, Wolf YI: Evolutionary Genomics of Defense Systems in Archaea and Bacteria. Annu Rev Microbiol 2017, 71:233–261.

106. Hussain FA, Dubert J, Elsherbini J, Murphy M, VanInsberghe D, Arevalo P, Kauffman K, Rodino-Janeiro BK, Gavin H, Gomez A, et al: Rapid evolutionary turnover of mobile genetic elements drives bacterial resistance to phages. Science 2021, 374:488–492.

107. Patel PH, Maxwell KL: Prophages provide a rich source of antiphage defense systems. Curr Opin Microbiol 2023, 73:102321.

108. Piel D, Bruto M, Labreuche Y, Blanquart F, Goudenege D, Barcia-Cruz R, Chenivesse S, Le Panse S, James A, Dubert J, et al: Phage-host coevolution in natural populations. Nat Microbiol 2022, 7:1075–1086.

109. Box AM, McGuffie MJ, O’Hara BJ, Seed KD: Functional Analysis of Bacteriophage Immunity through a Type I-E CRISPR-Cas System in *Vibrio cholerae* and Its Application in Bacteriophage Genome Engineering. J Bacteriol 2016, 198:578–590.

110. McDonald ND, Regmi A, Morreale DP, Borowski JD, Boyd EF: CRISPR-Cas systems are present predominantly on mobile genetic elements in *Vibrio* species. BMC Genomics 2019, 20:105.

111. Darracq B, Littner E, Brunie M, Bos J, Kaminski P-A, Depardieu F, Slesak W, Debatisse K, Touchon M, Bernheim A, et al: Sedentary chromosomal integrons as biobanks of bacterial anti-phage defence systems. bioRxiv 2024:2024.2007.2002.601686.

