## Supplemental material for "Exploring Mobile Genetic Elements in *Vibrio cholerae*"

#### SUPPLEMENTARY TABLES

**Table S1. Genome sequences of pandemic *V. cholerae* strains used in this study.**

| Strain | Description | Place of isolation | Year of isolation | GenBank accession numbers | Ref. |
|---|---|---|---|---|---|
| **MAK757** | O1 El Tor Ogawa, pre-pandemic | Celebes Island, Indonesia | 1937 | CP159790 / CP159791 | [1]; this study |
| **N16961** | O1 El Tor Inaba | Bangladesh | 1975 | CP028827 / CP028828 | [2, 3] |
| **E7946** | O1 El Tor Ogawa | Bahrain | 1978 | CP047303 / CP047304 | [4, 5] |
| **P27459** | O1 El Tor Inaba | Bangladesh | 1976 | CP047299 / CP047300 | [5, 6] |
| **A1552** | O1 El Tor Inaba | USA (Peruvian outbreak strain) | 1991 | CP028894 / CP028895 | [3, 7] |
| **C6706** | O1 El Tor Inaba | Peru | 1991 | CP047295 / CP047296 | [5, 8] |
| **DRC193A** | O1 El Tor | Democratic Republic of the Congo | 2011 | CP132180 / CP132181 | [9] |
| **DRC052** | O1 El Tor | Democratic Republic of the Congo | 2012 | CP132187 / CP132188 | [9] |
| **DRC072** | O1 El Tor | Democratic Republic of the Congo | 2009 | CP132182 / CP132183 | [9] |

**Table S2. Genome sequences of non-pandemic *V. cholerae* strains used in this study.**

| Strain | Description | Place of isolation | Year of isolation | GenBank accession numbers* | Ref. |
|---|---|---|---|---|---|
| **NCTC 30** | Patient isolate (British soldier), diarrhea; probably O2 serogroup | Egypt | 1916 | LS997867 / LS997868 | [10] |
| **VCSRO5 (RIMD2214243)** | Patient isolate (diarrhea); O5 serogroup | Philippines | 1964 | AP023377 / AP023378 | [11] |
| **VCSRO17 (RIMD2214255)** | Patient isolate (diarrhea); O17 serogroup | India | 1968 | AP023371 / AP023372 | [11] |
| **VCSRO45 (RIMD2214283)** | Patient isolate (diarrhea); O45 serogroup | India | 1973 | AP023375 / AP023376 | [11] |
| **VCSRO51 (RIMD2214289)** | Patient isolate (diarrhea); O51 serogroup | India | 1973 | AP023379 / AP023380 | [11] |
| **VCSRO63 (RIMD2214301)** | Patient isolate (diarrhea); O63 serogroup | India | 1976 | AP023381 / AP023382 | [11] |
| **VCSRO77 (RIMD2214315)** | Patient isolate (diarrhea); O77 serogroup | India | 1976 | AP023383 / AP023384 | [11] |
| **VCSRO96 (RIMD2214334)** | Patient isolate (diarrhea); O96 serogroup | India | 1976 | AP02338 / AP023386 | [11] |
| **VCSRO102 (RIMD2214340)** | Patient isolate (diarrhea); O102 serogroup | China | 1988 | AP023331 / AP023332 | [11] |
| **VCSRO162 (RIMD2214400)** | Patient isolate (diarrhea); O162 serogroup | Argentina | 1993 | AP023369 / AP023370 | [11] |
| **SIO** | Environmental isolate | Scripps Institute of Oceanography Pier (CA, USA) | 2000 | JBBHLM000000000.1 | [12, 13] |
| **TP** | Environmental isolate (plankton) | Torrey Pines Estuary (CA, USA) | 2000 | CP137095 / CP137096 | [12, 13] |
| **VCSRO207 (RIMD2214445)** | Environmental isolate; O207 serogroup | Japan | 2001 | AP023373 / AP023374 | [11] |
| **E7G** | Environmental isolate | Moss Landing Harbor (CA, USA) | 2004 | CP053822 / CP053823 | [14, 15] |
| **L6G** | Environmental isolate | Lagunitas Creek (CA, USA) | 2004 | CP053802 / CP053803 | [14, 15] |
| **SA10G** | Environmental isolate | Old Salinas River (CA, USA) | 2004 | CP053820 / CP053821 | [14, 15] |
| **SA3G** | Environmental isolate | Old Salinas River (CA, USA) | 2004 | CP053744 / CP053745 | [14, 15] |

| | | | | | |
|---|---|---|---|---|---|
| **SA5Y** | Environmental isolate | Old Salinas River (CA, USA) | 2004 | CP028892 / CP028893 | [14, 15] |
| **SA7G** | Environmental isolate | Old Salinas River (CA, USA) | 2004 | CP053816 / CP053817 | [14, 15] |
| **SL4G** | Environmental isolate | San Lorenzo River (CA, USA | 2004 | CP053796 / CP053797 | [14, 15] |
| **SL5Y** | Environmental isolate | San Lorenzo River (CA, USA) | 2004 | CP053798 / CP053799 | [14, 15] |
| **SL6Y** | Environmental isolate | San Lorenzo River (CA, USA) | 2004 | CP053804 / CP053805 | [14, 15] |
| **SO5Y** | Environmental isolate | Soquel Creek (CA, USA) | 2004 | CP053800 / CP053801 | [14, 15] |
| **SP6G** | Environmental isolate | San Pedro Creek (CA, USA) | 2004 | CP053806 / CP053807 | [14, 15] |
| **SP7G** | Environmental isolate | San Pedro Creek (CA, USA) | 2004 | CP053808 / CP053809 | [14, 15] |
| **W10G** | Environmental isolate | Waddell Creek (CA, USA) | 2004 | CP053794 / CP053795 | [14, 15] |
| **W6G** | Environmental isolate | Waddell Creek (CA, USA) | 2004 | CP053810 / CP053811 | [14, 15] |
| **W7G** | Environmental isolate | Waddell Creek (CA, USA) | 2004 | CP053813 / CP053814 | [14, 15] |
| **DL4211** | Environmental isolate, O123 serogroup | Rio Grande river (USA) | 2008 | CP137091 / CP137092 | [13, 16] |
| **DRC187** | Non-O1 isolate; *ctx*- and *tcp*-negative | Sud Kivu, Democratic Republic of the Congo | 2011 | CP138190 / CP132189 | [9] |
| **2012Env-390** | Environmental isolate; non-toxigenic O1 Ogawa | Gressier, Haiti | 2012 | CP013013 / CP013014 | [17, 18] |
| **2012Env-9** | Environmental isolate; non-toxigenic O1 Ogawa | La Salle, Haiti | 2012 | CP012997 / CP012998 | [17, 18] |
| **V130003** | Patient isolate (feces); O144 serogroup | Japan | 2014 | AP024967 / AP024968 | - |
| **Strain 20000** | Environmental isolate; O1 serogroup | Rostov-on-Don, Russia | 2016 | CP036499 / CP036500 | - |
| **RFB05** | Environmental isolate | North Park Lake (PA, USA) | 2017 | CP043557 / CP043558 | - |
| **RFB16** | Environmental isolate | North Park Lake (PA, USA) | 2017 | CP043554 / CP043556 | [19] |

* If two accession numbers are provided, they represent chromosomes 1 and 2.

**Table S3. Software tools used to identify genomic islands in each genome.**

| Program | Version | Reference |
|---|---|---|
| **Alienhunter** | 1.7 | [20] |
| **IslandCafe** | 1.0 | [21] |
| **Colombo: SigiHMM** | 4.0 | [22] |
| **Colombo: SigiCRF** | 4.0 | [22] |
| **IslandPath-DIMOB** | December 2016 | [23, 24] |

**Table S4. Genome sequences of *Vibrios* and *Photobacterium* species used in this study.**

| Species | Strain | Place of isolation | Year of isolation | GenBank accession numbers* | Ref. |
|---|---|---|---|---|---|
| ***Photobacterium gaetbulicola*** | Gung 47 | Korea | N/A | CP005973.1 / CP005974.1 | - |
| ***Vibrio alginolyticus*** | E110 | South China | 2019 | CP098033.1 / CP098034.1 | [25] |
| ***Vibrio alginolyticus*** | ATCC17749 | China | 2015 | CP006718.1 / CP006719.1 | [26] |
| ***Vibrio furnissii*** | FDAARGOS_777 | USA | N/A | CP040990.1 / CP040991.1 | - |
| ***Vibrio furnissii*** | RM8376 | N/A | N/A | CP064379.1 / CP064381.1 | - |
| ***Vibrio mimicus*** | ATCC33655 | N/A | 2017 | NZ_LOSJ01000001 / NZ_LOSJ01000002 | - |
| ***Vibrio mimicus*** | SCCF01 | China | 2013 | CP016383.1 / CP016384.1 | - |
| ***Vibrio tarriae*** | 2521-89 | USA | 1989 | CP022353.1 / CP022352.1 | - |
| ***Vibrio vulnificus*** | ATCC27562 | USA | 1979 | CP012881.1 / CP012882.1 | [27] |
| ***Vibrio vulnificus*** | FORC037 | South Korea | 1993 | CP016321.1 / CP016322.1 | - |

* If two accession numbers are provided, they represent chromosomes 1 and 2.

**Table S5. Insertion sites in chromosome 1.**

| | Insertion site | Locus tag[1] | Details for non-coding insertion sites |
|---|---|---|---|
| 1 | ***mnmE*** | *VC0003* | |
| 2 | ***gyrB*** | *VC0015* | |
| 3 | ***comM*** | *VC0032* | |
| 4 | ***rmuC*** | *VC0082* | |
| 5 | ***yijD*** | *VC0153* | |
| 6 | ***gor*** | *VC0186* | |
| 7 | ***rph*** | *VC0210* | |
| 8 | **O-antigen** | *VC0264* | Locus tag for *rjg* gene, next to O-antigen |
| 9 | ***dus*** | *VC0291* | |
| 10 | ***yjbQ*** | *VC0373* | |
| 11 | ***csrD*** | *VC0398* | |
| 12 | ***tRNA-Met*** | *VC0517* | Locus tag for *rpoD* gene, next to *tRNA-Met* (tRNA annotated as tRNA-Met by Heidelberg *et al.* [2]; *tRNA* prediction tools suggest locus *tRNA-Ile*). |
| 13 | ***prfC*** | *VC0659* | |
| 14 | ***guaA*** | *VC0768* | |
| 15 | ***ssrA*** | *VC0848* | Locus tag for *smpB* gene, next to *ssrA tmRNA* |
| 16 | ***tRNA-Asn*** | *VC1018* | Locus tag for *uvrB* gene, next to *tRNA-Asn* |
| 17 | ***ffs*** | *VC1067* | Locus tag for *yeaP* gene, next to *ffs ncRNA* |
| 18 | ***sbcB*** | *VC1234* | |
| 19 | ***rtxA*** | *VC1451* | |
| 20 | ***pepN*** | *VC1494* | |
| 21 | ***katG*** | *VC1560* | |
| 22 | ***tRNA-Val*** | *VC1598* | Locus tag for hypothetical gene, next to *tRNA-Val* |
| 23 | ***bepC*** | *VC1621* | |
| 24 | ***tRNA-Ser*** | *VC1757* | Locus tag for *swrC* gene, next to *tRNA-Ser* |
| 25 | ***pflB*** | *VC1866* | |
| 26 | ***pyrF*** | *VC1911* | |
| 27 | ***tRNA-Arg*** | *VC1942* | Locus tag for *folD* gene, next to *tRNA-Arg* |
| 28 | ***rlmA*** | *VC2003* | |
| 29 | ***tRNA-Pro*** | *VC2041* | Locus tag for *yejM* gene, next to *tRNA-Pro* |
| 30 | ***topB*** | *VC2043* | |
| 31 | ***tRNA-Leu*** | *VC2187* | Locus tag for *flaC* gene, next to *tRNA-Leu* |
| 32 | ***thrC*** | *VC2362* | |
| 33 | ***VC2384*** | *VC2384* | |

[1] Locus tags are according to reference genome of strain N16961 [2].

**Table S6. Insertion sites in chromosome 2.**

| | Insertion site | Locus tag[1] | Details for non-coding insertion sites |
|---|---|---|---|
| 1 | ***guaC*** | *VCA0197* | |
| 2 | ***ackA*** | *VCA0235* | |
| 3 | ***gcvT*** | *VCA0280* | |
| 4 | **Integron** | *VCA0291* | Integron integrase gene |
| 5 | ***nrdD*** | *VCA0511* | |
| 6 | ***clcA*** | *VCA0526* | |
| 7 | ***pntB*** | *VCA0564* | |
| 8 | ***nfsb*** | *VCA0637* | |
| 9 | *VCA0735* | *VCA0735* | Annotated as encoding a hypothetical protein |
| 10 | ***ugd*** | *VCA0780* | |
| 11 | ***yjdF*** | *VCA0789* | |
| 12 | ***tadA*** | *VCA0840* | |
| 13 | ***tdh*** | *VCA0885* | |
| 14 | ***glgX*** | *VCA1029* | |

[1] Locus tags are according to reference genome of strain N16961 [2].

**Table S7. Locus tags corresponding to *rjg* gene (*VC0264* in N16961[1]) in non-pandemic strains.**

| Strain | *rjg* locus tag |
|---|---|
| **NCTC 30** | SAMEA104470976_02470 |
| **VCSRO5 (RIMD2214243)** | VCSRO5_2605 |
| **VCSRO17 (RIMD2214255)** | VCSRO17_0348 |
| **VCSRO45 (RIMD2214283)** | VCSRO45_2616 |
| **VCSRO51 (RIMD2214289)** | VCSRO51_2578 |
| **VCSRO63 (RIMD2214301)** | VCSRO63_2503 |
| **VCSRO77 (RIMD2214315)** | VCSRO77_0347 |
| **VCSRO96 (RIMD2214334)** | VCSRO96_2490 |
| **VCSRO102 (RIMD2214340)** | VCSRO102_2484 |
| **VCSRO162 (RIMD2214400)** | VCSRO162_2560 |
| **SIO** | R4537_17560 |
| **TP** | R4536_06825 |
| **VCSRO207 (RIMD2214445)** | VCSRO207_2465 |
| **E7G** | HPY10_12585 |
| **L6G** | HPY16_12095 |
| **SA10G** | HPY11_12655 |
| **SA3G** | HPY04_12800 |
| **SA5Y** | Sa5Y_VC02581 |
| **SA7G** | HPY17_12545 |
| **SL4G** | HPY13_12845 |
| **SL5Y** | HPY14_12725 |
| **SL6Y** | HPY06_12480 |
| **SO5Y** | HPY15_12665 |
| **SP6G** | HPY07_12630 |
| **SP7G** | HPY08_12200 |
| **W10G** | HPY12_13210 |
| **W6G** | HPY05_12810 |
| **W7G** | HPY09_12815 |
| **DL4211** | R4535_00585 |
| **DRC-187A** | Q9L40_10905 |
| **2012Env-390** | AP033_RS17895 |
| **2012Env-9** | NH62_RS17080 |
| **V130003** | V130003_25610 |
| **Strain 20000** | EZZ25_03155 |
| **RFB05** | F0315_04125 |
| **RFB16** | F0316_12775 |

[1] Locus tags are according to reference genome of strain N16961 [2].

#### SUPPLEMENTARY REFERENCES


1. Chattopadhyay DJ, Sarkar BL, Ansari MQ, Chakrabarti BK, Roy MK, Ghosh AN, Pal SC: **New phage typing scheme for *Vibrio cholerae* O1 biotype El Tor strains.** *J Clin Microbiol* 1993, **31:**1579-1585.
2. Heidelberg JF, Eisen JA, Nelson WC, Clayton RA, Gwinn ML, Dodson RJ, Haft DH, Hickey EK, Peterson JD, Umayam L, et al: **DNA sequence of both chromosomes of the cholera pathogen *Vibrio cholerae*.** *Nature* 2000, **406:**477-483.
3. Matthey N, Drebes Dörr NC, Blokesch M: **Long-Read-Based Genome Sequences of Pandemic and Environmental *Vibrio cholerae* Strains.** *Microbiol Resour Announc* 2018, **7:**e01574-01518.
4. Miller VL, DiRita VJ, Mekalanos JJ: **Identification of *toxS*, a regulatory gene whose product enhances ToxR-mediated activation of the cholera toxin promoter.** *J Bacteriol* 1989, **171:**1288-1293.
5. Stutzmann S, Blokesch M: **Comparison of chitin-induced natural transformation in pandemic *Vibrio cholerae* O1 El Tor strains.** *Environ Microbiol* 2020, **22:**4149-4166.
6. Pearson GD, Woods A, Chiang SL, Mekalanos JJ: **CTX genetic element encodes a site-specific recombination system and an intestinal colonization factor.** *Proc Natl Acad Sci USA* 1993, **90:**3750-3754.
7. Yildiz FH, Schoolnik GK: **Role of *rpoS* in stress survival and virulence of *Vibrio cholerae*.** *J Bacteriol* 1998, **180:**773-784.
8. Stutzmann S, Blokesch M: **Circulation of a Quorum-Sensing-Impaired Variant of *Vibrio cholerae* Strain C6706 Masks Important Phenotypes.** *mSphere* 2016, **1:e00098-16**.
9. Lemopoulos A, Miwanda B, Drebes Dörr Natália C, Stutzmann S, Bompangue D, Muyembe-Tamfum J-J, Blokesch M: **Genome sequences of Vibrio cholerae strains isolated in the DRC between 2009 and 2012.** *Microbio Resour Announc* 2024, **0:**e00827-00823.
10. Dorman MJ, Kane L, Domman D, Turnbull JD, Cormie C, Fazal MA, Goulding DA, Russell JE, Alexander S, Thomson NR: **The history, genome and biology of NCTC 30: a non-pandemic Vibrio cholerae isolate from World War One.** *Proc Biol Sci* 2019, **286:**20182025.
11. Murase K, Arakawa E, Izumiya H, Iguchi A, Takemura T, Kikuchi T, Nakagawa I, Thomson NR, Ohnishi M, Morita M: **Genomic dissection of the *Vibrio cholerae* O-serogroup global reference strains: reassessing our view of diversity and plasticity between two chromosomes.** *Microb Genom* 2022, **8**.
12. Purdy A, Rohwer F, Edwards R, Azam F, Bartlett DH: **A glimpse into the expanded genome content of Vibrio cholerae through identification of genes present in environmental strains.** *J Bacteriol* 2005, **187:**2992-3001.

13. Otto SB, Servajean R, Lemopoulos A, Bitbol AF, Blokesch M: **Interactions between pili affect the outcome of bacterial competition driven by the type VI secretion system.** *Curr Biol* 2024, **34:**2403-2417 e2409.
14. Keymer DP, Miller MC, Schoolnik GK, Boehm AB: **Genomic and phenotypic diversity of coastal *Vibrio cholerae* strains is linked to environmental factors.** *Appl Environ Microbiol* 2007, **73:**3705-3714.
15. Drebes Dörr NC, Blokesch M: **Interbacterial competition and anti-predatory behavior of environmental *Vibrio cholerae* strains.** *Environ Microbiol* 2020, **22:**4485-4504.
16. Unterweger D, Kitaoka M, Miyata ST, Bachmann V, Brooks TM, Moloney J, Sosa O, Silva D, Duran-Gonzalez J, Provenzano D, Pukatzki S: **Constitutive type VI secretion system expression gives *Vibrio cholerae* intra- and interspecific competitive advantages.** *PLoS One* 2012, **7:**e48320.
17. Azarian T, Ali A, Johnson JA, Mohr D, Prosperi M, Veras NM, Jubair M, Strickland SL, Rashid MH, Alam MT, et al: **Phylodynamic analysis of clinical and environmental *Vibrio cholerae* isolates from Haiti reveals diversification driven by positive selection.** *mBio* 2014, **5**.
18. Azarian T, Ali A, Johnson JA, Jubair M, Cella E, Ciccozzi M, Nolan DJ, Farmerie W, Rashid MH, Sinha-Ray S, et al: **Non-toxigenic environmental *Vibrio cholerae* O1 strain from Haiti provides evidence of pre-pandemic cholera in Hispaniola.** *Sci Rep* 2016, **6:**36115.
19. Bina RF, Bina JE, Weng Y: **Genome Sequence of *Vibrio cholerae* Strain RFB16, Isolated from North Park Lake in Allegheny County, Pennsylvania.** *Microbiol Resour Announc* 2020, **9**.
20. Vernikos GS, Parkhill J: **Interpolated variable order motifs for identification of horizontally acquired DNA: revisiting the Salmonella pathogenicity islands.** *Bioinformatics* 2006, **22:**2196-2203.
21. Jani M, Azad RK: **IslandCafe: Compositional Anomaly and Feature Enrichment Assessment for Delineation of Genomic Islands.** *G3 (Bethesda)* 2019, **9:**3273-3285.
22. Waack S, Keller O, Asper R, Brodag T, Damm C, Fricke WF, Surovcik K, Meinicke P, Merkl R: **Score-based prediction of genomic islands in prokaryotic genomes using hidden Markov models.** *BMC Bioinformatics* 2006, **7:**142.
23. Bertelli C, Brinkman FSL: **Improved genomic island predictions with IslandPath-DIMOB.** *Bioinformatics* 2018, **34:**2161-2167.
24. Hsiao WW, Ung K, Aeschliman D, Bryan J, Finlay BB, Brinkman FS: **Evidence of a large novel gene pool associated with prokaryotic genomic islands.** *PLoS Genet* 2005, **1:**e62.
25. Li X, Zhang C, Jin X, Wei F, Yu F, Call DR, Zhao Z: **Temporal Transcriptional Responses of a *Vibrio alginolyticus* Strain to Podoviridae Phage HH109 Revealed by RNA-Seq.** *mSystems* 2022, **7:**e0010622.

26. Liu XF, Cao Y, Zhang HL, Chen YJ, Hu CJ: **Complete Genome Sequence of *Vibrio alginolyticus* ATCC 17749T.** *Genome Announc* 2015, **3**.
27. Rusch DB, Rowe-Magnus DA: **Complete Genome Sequence of the Pathogenic *Vibrio vulnificus* Type Strain ATCC 27562.** *Genome Announc* 2017, **5**.

#### SUPPLEMENTARY FIGURE LEGENDS & FIGURES

**Figure S1. Conservation of MGEs in pandemic *V. cholerae* strains (part I). a)** Phylogenetic analysis of *Vibrio* species. This figure showcases the evolutionary relationship between selected *V. cholerae* strains from this study and other *Vibrio* and *Photobacterium* species (Table S4). Notably, strain RFB05, previously classified as *V. cholerae*, is identified as belonging to the *V. tarriae* species, highlighted by the red box. The phylogenetic tree was recreated using IQ-TREE2 (v.2.2.0). The core protein sequences (457 total proteins, with the --phylo parameter) of the species were concatenated, derived from the PPanGGOLiN (v.1.2.74) pipeline (fasta and annotation files provided, with the rest of the parameters set to default). **b-d)** Gene content comparison of pandemic islands. These panels display the comparative gene content of key pandemic islands VPI-2 **(b),** and VSP-I **(c),** and VSP-II **(d)** across various strains. Light grey connections signify protein identities above 30% (default parameter of the clinker pipeline). These connections are not present in the case of tRNA genes such as *tRNA-Ser* **(b)** and *tRNA-Met* **(d)**, as comparisons in clinker are based on protein sequences. Insertion sites are marked in green (arrows in the case of protein-coding genes, small squares in the case of tRNA genes). Dark grey arrows point to genes flanking the islands, with important genes labelled. Light grey arrows denote genes encoding hypothetical proteins or those not directly related to the main findings. Light blue arrows highlight mobilization-related genes. Yellow arrows mark antiviral defense genes or gene clusters, while salmon-colored arrows indicate phage defence candidates (PDC), each labelled accordingly. The gene marked with “WYL*” was identified by a loose model that detects various defense-modification systems (“Dms_other”) by PADLOC. The *nan-nag* region in VPI-2 is marked in purple. **e)** Nucleotide sequence-based alignment heatmap for VSP-I of strains DRC072 and DRC052, carried on chromosomes 1 or 2.

**Figure S2. Conservation of MGEs in pandemic *V. cholerae* strains (part II). a-d, g-i)** Nucleotide sequence alignment-based heatmaps illustrating the genetic similarity of islands identified across various pandemic *V. cholerae* strains. Specific genomic islands are VPI-1 and VPI-2 (panels **a** and **b**), VSP-I and II (panels **c** and **d**), Vc95 Retron **(g)**, and islands inserted at *guaC* **(h)** and *tdh* **(i)**. **e-f)** Gene content comparison of pandemic islands WASA-1 **(e)** and ICE*Vch*Ind5 **(f)**. Light grey connections show areas where protein identity exceeds 30%, following default analysis settings in Clinker. Insertion site genes (*pepN* and *prfC*) are marked in green. Color code as in Fig. S1.

**Figure S3. Comparative gene content of genomic islands at the *gcvT* locus of non-pandemic *V. cholerae***. This figure presents the gene content within genomic islands located at the *gcvT* locus (marked in green) across different non-pandemic *V. cholerae* strains, listed on the left side (reduced size island found in A1552 kept for comparison). Light grey connections show areas where protein identity exceeds 30%, following default analysis settings of the clinker pipeline. Color code as in Fig. S1. T6SS Aux3 clusters are highlighted in dark pink.

**Figure S4. Comparative gene content of genomic islands at the *ssrA* locus of non-pandemic *V. cholerae***. This figure presents the gene content within genomic islands located at the *ssrA* locus (marked in green) across different non-pandemic *V. cholerae* strains, listed on the left side (VPI-1 island found in A1552 was kept for comparison). Light grey connections show areas where protein identity exceeds 30%, following default analysis settings of clinker. These connections are not present in the case of *ssrA* specifically, as it is a tmRNA, and connections in clinker are based on protein-coding sequences only. Color code as in Fig. S1. The presence of TCP + Acf clusters, akin to those in the pandemic VPI-1 island, is noted in light pink, whereas T6SS Aux4 clusters are distinguished in dark pink.

**Figure S5. Comparative gene content of genomic islands found at the *tdh* locus of non-pandemic *V. cholerae*.** Strains harboring the specified genomic islands inserted at *tdh* (marked in green) are indicated on the left. Color code as for Figs. S1, with T6SS Aux5 clusters marked in pink. An operon related to $Ni^{2+}$ and $K^{+}$ transport, which is also observed in other location (see Fig. S6), is marked in red. Genes potentially related to antiviral defense systems are labelled on the 3' ends of the islands.

**Figure S6. Comparative gene content of genomic islands found at the *pntB* locus of non-pandemic *V. cholerae*.** Strains harboring the specified genomic islands inserted at *pntB* (marked in green) are indicated on the left. Color code as for Figs. S1 and S5. T6SS Aux5 clusters are distinguished in dark pink.

**Figure S7. Comparative gene content of genomic islands found at the *VC2384* locus of non-pandemic *V. cholerae*.** Genomic islands inserted at *VC2384* (marked in green) are indicated with their host strains indicated on the left. Color code as for Fig. S1. The island found in pandemic strain A1552 (Vc95 Retron) is kept for comparison. Genes potentially related to antiviral defense systems are labelled.

**Figure S8. Comparative gene content of genomic islands found at the *ffs* locus of non-pandemic *V. cholerae*.** Genomic islands inserted at *ffs* (ncRNA marked with a small green rectangle) are indicated with their host strains indicated on the left. Light grey connections show areas where protein identity exceeds 30%, following default analysis settings. These connections are not present in the case of *ffs* specifically, as it is a ncRNA, and connections are based on protein-coding sequences only. Color code as for Fig. S1.

**Figure S9. Comparative gene content of genomic islands found at the *rph* and *mnmE* loci of non-pandemic *V. cholerae*.** Genomic islands inserted at *rph* **(a)** and *mnmE* **(b)** are indicated with their host strains indicated on the left. Insertion site genes are marked in green. Color code as for Fig. S1.

**Figure S10. Comparative gene content of genomic islands found at the *guaC* locus of non-pandemic *V. cholerae*.** Genomic islands inserted at *guaC* (marked in green) are indicated with their host strains indicated on the left. Color code as for Figs. S1 and S5. The island found in pandemic strain A1552 (Menshen defense system) is kept for comparison. In islands found in DRC187 and A1552, the RM type II system genes are marked in faint orange to differentiate them from the Menshen defense system (in yellow).

**Figure S11. Comparative gene content of genomic islands found in *VCA0735* in non-pandemic *V. cholerae*.** Strains harboring the specified genomic islands inserted at *VCA0735* (marked in green) are indicated on the left. Color code as for Fig. S1. A piscibactin siderophore biosynthesis cluster and the *betTIBA* cluster related to osmoprotection are indicated. Genes potentially related to antiviral defense systems are labelled on the 3’ ends of the islands.

**Figure S12. Comparative gene content of genomic islands found at the *yijD* and *tRNA-Arg* loci of non-pandemic *V. cholerae*.** Strains harboring the specified genomic islands inserted at *yijD* **(a)** and *tRNA-Arg* **(b)** are indicated on the left. Insertion site protein-coding genes and tRNA genes are marked in green. Light grey connections show areas where protein identity exceeds 30%, following default analysis settings. These connections are not present in the case of *tRNA-Arg* specifically, as it is a tRNA, and connections are based on protein-coding sequences only. Color code as for Fig. S1 and S5.

Figure S1

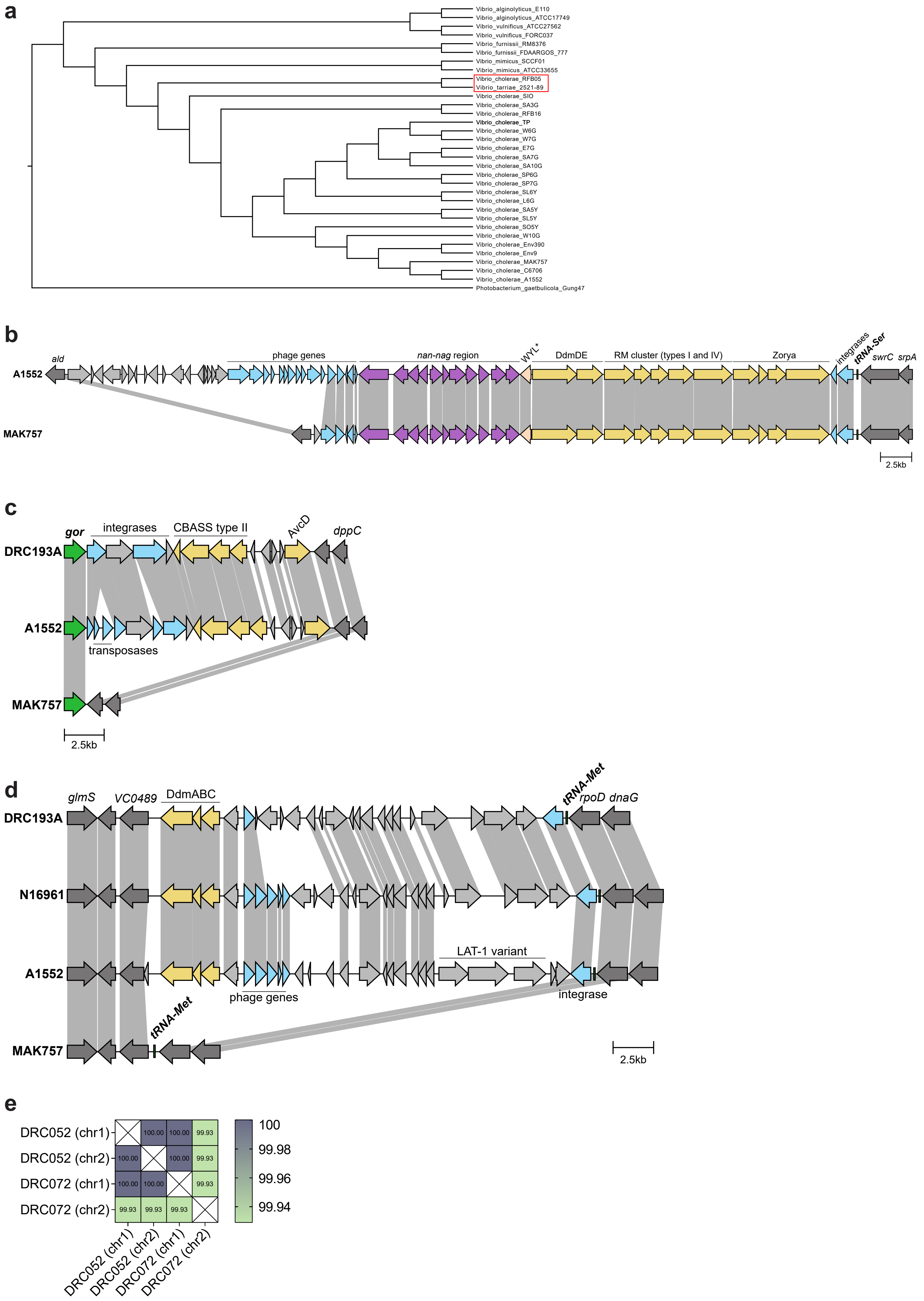
a
Vibrio_alginolyticus_E110
Vibrio_alginolyticus_ATCC17749
Vibrio_vulnificus_ATCC27562
Vibrio_vulnificus_FORC037
Vibrio_furnissii_RM8376
Vibrio_furnissii_FDAARGOS_777
Vibrio_mimicus_SCCF01
Vibrio_mimicus_ATCC33655
Vibrio_cholerae_RFB05
Vibrio_tarriae_2521-89
Vibrio_cholerae_SIO
Vibrio_cholerae_SA3G
Vibrio_cholerae_RFB16
Vibrio_cholerae_TP
Vibrio_cholerae_W6G
Vibrio_cholerae_W7G
Vibrio_cholerae_E7G
Vibrio_cholerae_SA7G
Vibrio_cholerae_SA10G
Vibrio_cholerae_SP6G
Vibrio_cholerae_SP7G
Vibrio_cholerae_SL6Y
Vibrio_cholerae_L6G
Vibrio_cholerae_SA5Y
Vibrio_cholerae_SL5Y
Vibrio_cholerae_SO5Y
Vibrio_cholerae_W10G
Vibrio_cholerae_Env390
Vibrio_cholerae_Env9
Vibrio_cholerae_MAK757
Vibrio_cholerae_C6706
Vibrio_cholerae_A1552
Photobacterium_gaetbulicola_Gung47
b
ald
phage genes
nan-nag region
WYL*
DdmDE
RM cluster (types I and IV)
Zorya
integrases
tRNA-Ser
swrC
srpA
A1552
MAK757
2.5kb
c
gor
integrases
CBASS type II
AvcD
dppC
DRC193A
A1552
transposases
MAK757
2.5kb
d
glmS
VC0489
DdmABC
tRNA-Met
rpoD
dnaG
DRC193A
N16961
LAT-1 variant
A1552
phage genes
integrase
MAK757
tRNA-Met
2.5kb
e
DRC052 (chr1)
DRC052 (chr2)
DRC072 (chr1)
DRC072 (chr2)
100.00
99.93
100
99.98
99.96
99.94

Figure S2

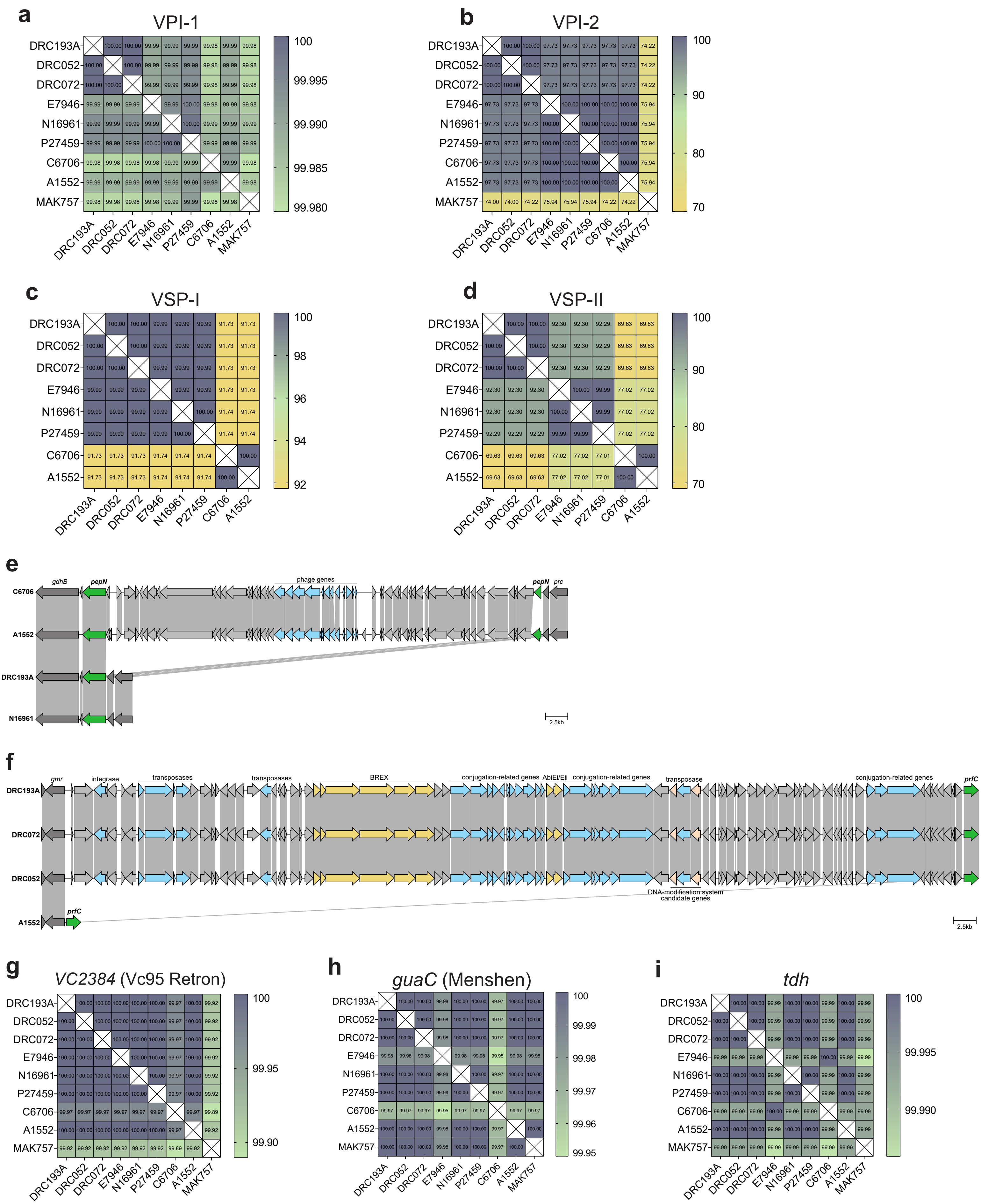
a
VPI-1
b
VPI-2
c
VSP-I
d
VSP-II
DRC193A
DRC052
DRC072
E7946
N16961
P27459
C6706
A1552
MAK757
e
gdhB
pepN
phage genes
prc
2.5kb
f
gmr
integrase
transposases
BREX
conjugation-related genes
AbiEi/Eii
transposase
prfC
DNA-modification system candidate genes
2.5kb
g
VC2384 (Vc95 Retron)
h
guaC (Menshen)
i
tdh

Figure S3

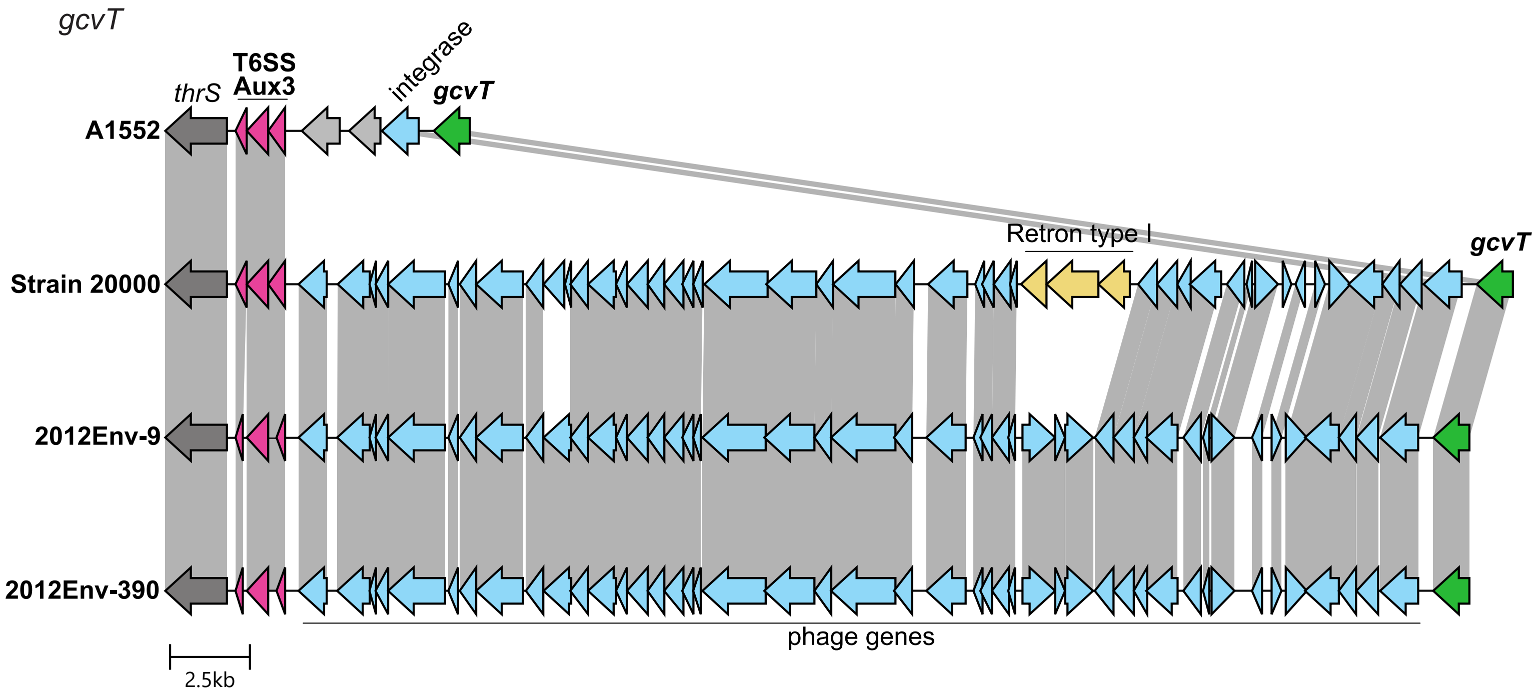
gcvT
T6SS
Aux3
thrS
integrase
gcvT
A1552
Retron type I
gcvT
Strain 20000
2012Env-9
2012Env-390
phage genes
2.5kb

Figure S4

*ssrA*

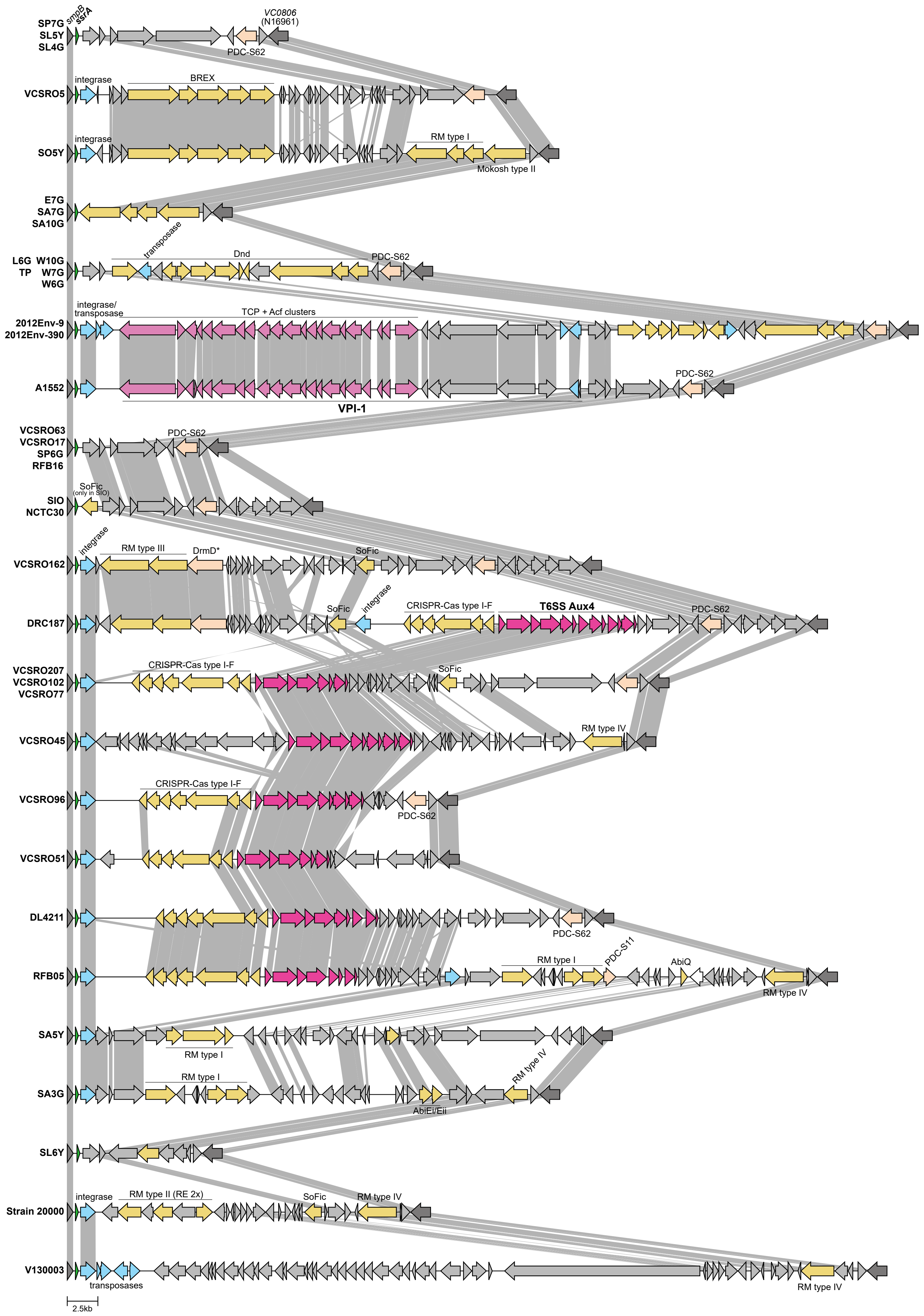

### Figure S5

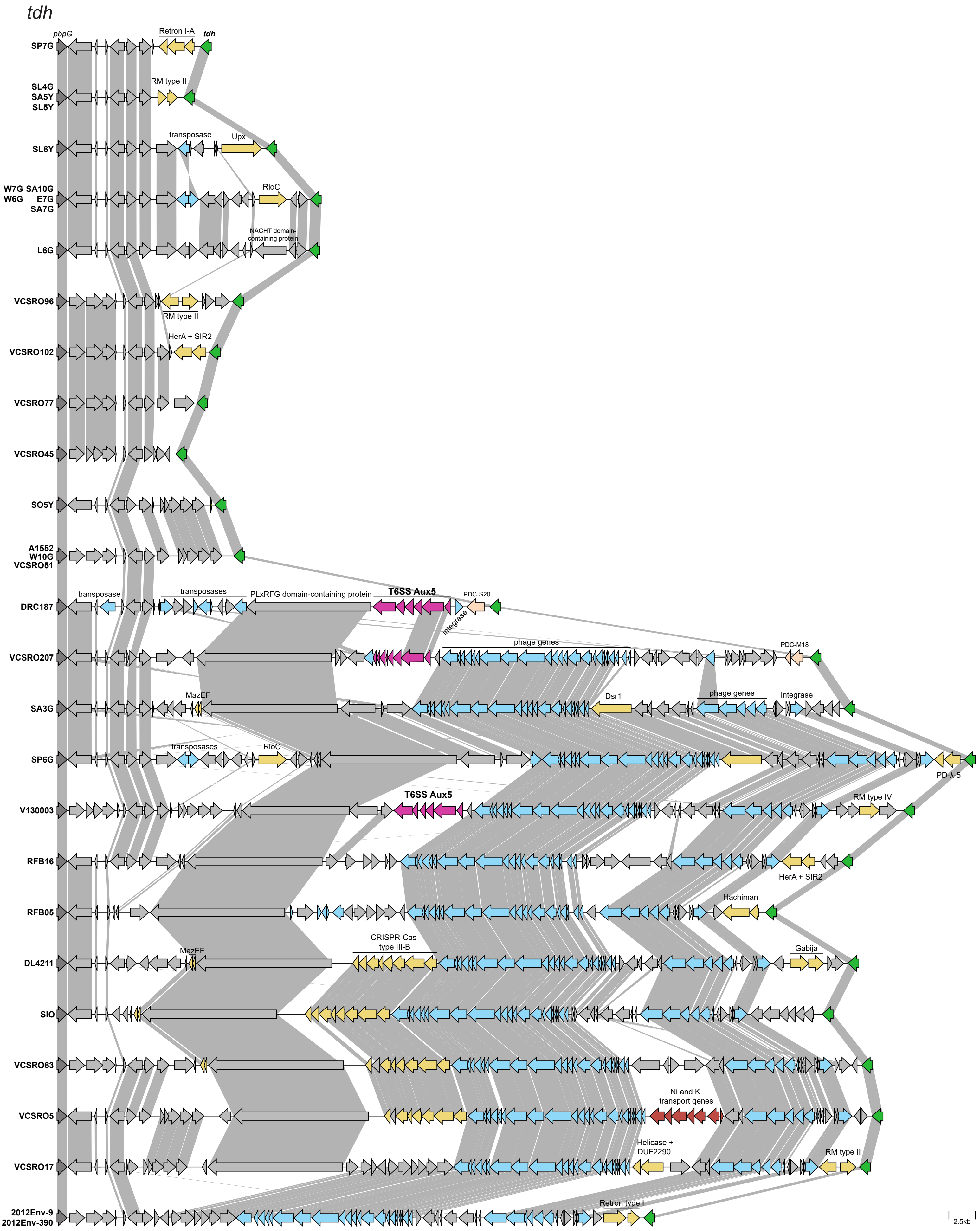
tdh
pbpG
Retron I-A
tdh
SP7G
SL4G
SA5Y
SL5Y
RM type II
SL6Y
transposase
Upx
W7G SA10G
W6G E7G
SA7G
RloC
L6G
NACHT domain-containing protein
VCSRO96
RM type II
VCSRO102
HerA + SIR2
VCSRO77
VCSRO45
SO5Y
A1552
W10G
VCSRO51
DRC187
transposase
transposases
PLxRFG domain-containing protein
T6SS Aux5
PDC-S20
integrase
VCSRO207
phage genes
PDC-M18
SA3G
MazEF
Dsr1
phage genes
integrase
SP6G
transposases
RloC
PD-λ-5
V130003
T6SS Aux5
RM type IV
RFB16
HerA + SIR2
RFB05
Hachiman
DL4211
MazEF
CRISPR-Cas
type III-B
Gabija
SIO
VCSRO63
VCSRO5
Ni and K
transport genes
VCSRO17
Helicase +
DUF2290
RM type II
2012Env-9
2012Env-390
Retron type I
2.5kb

Figure S6

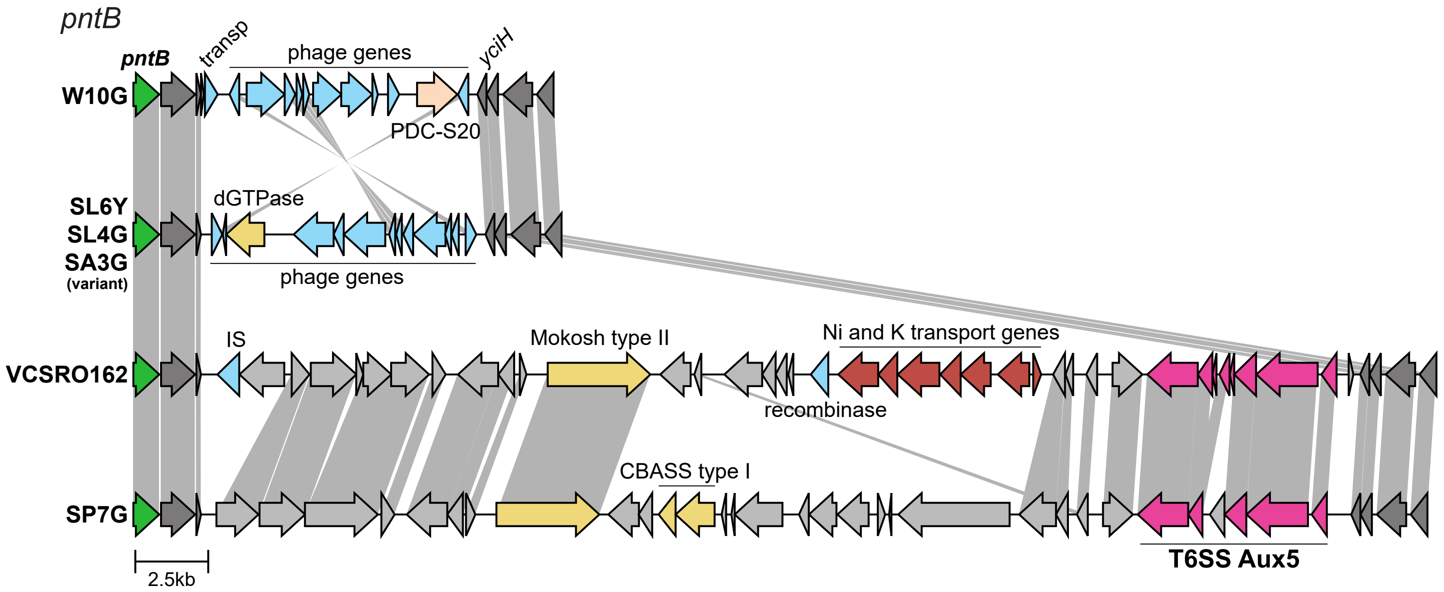

pntB
pntB
transp
phage genes
yciH
W10G
PDC-S20
dGTPase
SL6Y
SL4G
SA3G
(variant)
phage genes
IS
Mokosh type II
Ni and K transport genes
VCSRO162
recombinase
CBASS type I
SP7G
T6SS Aux5
2.5kb

### Figure S7

*VC2384*

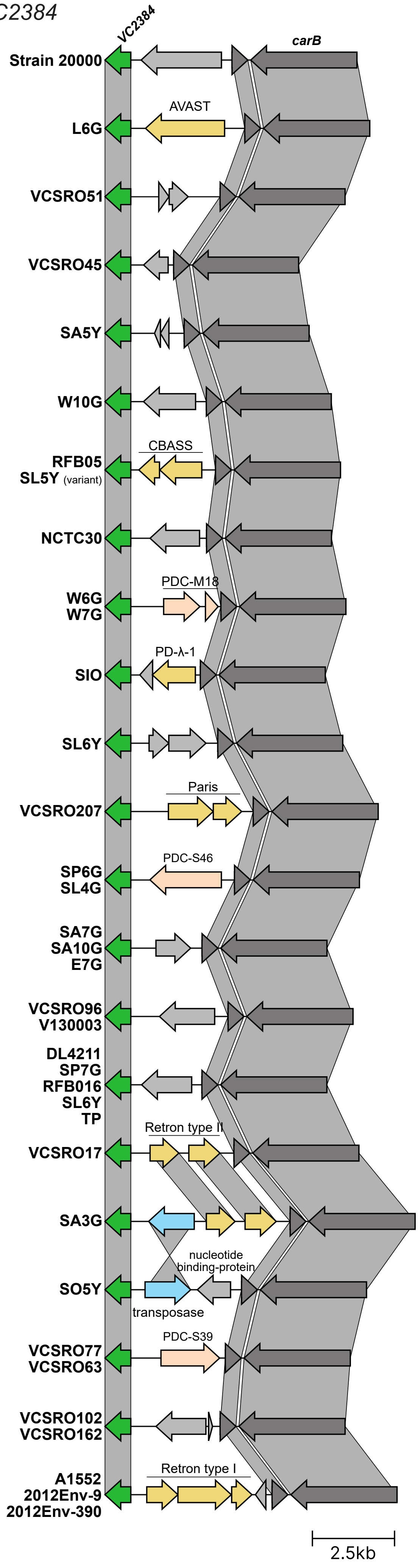

Figure S8

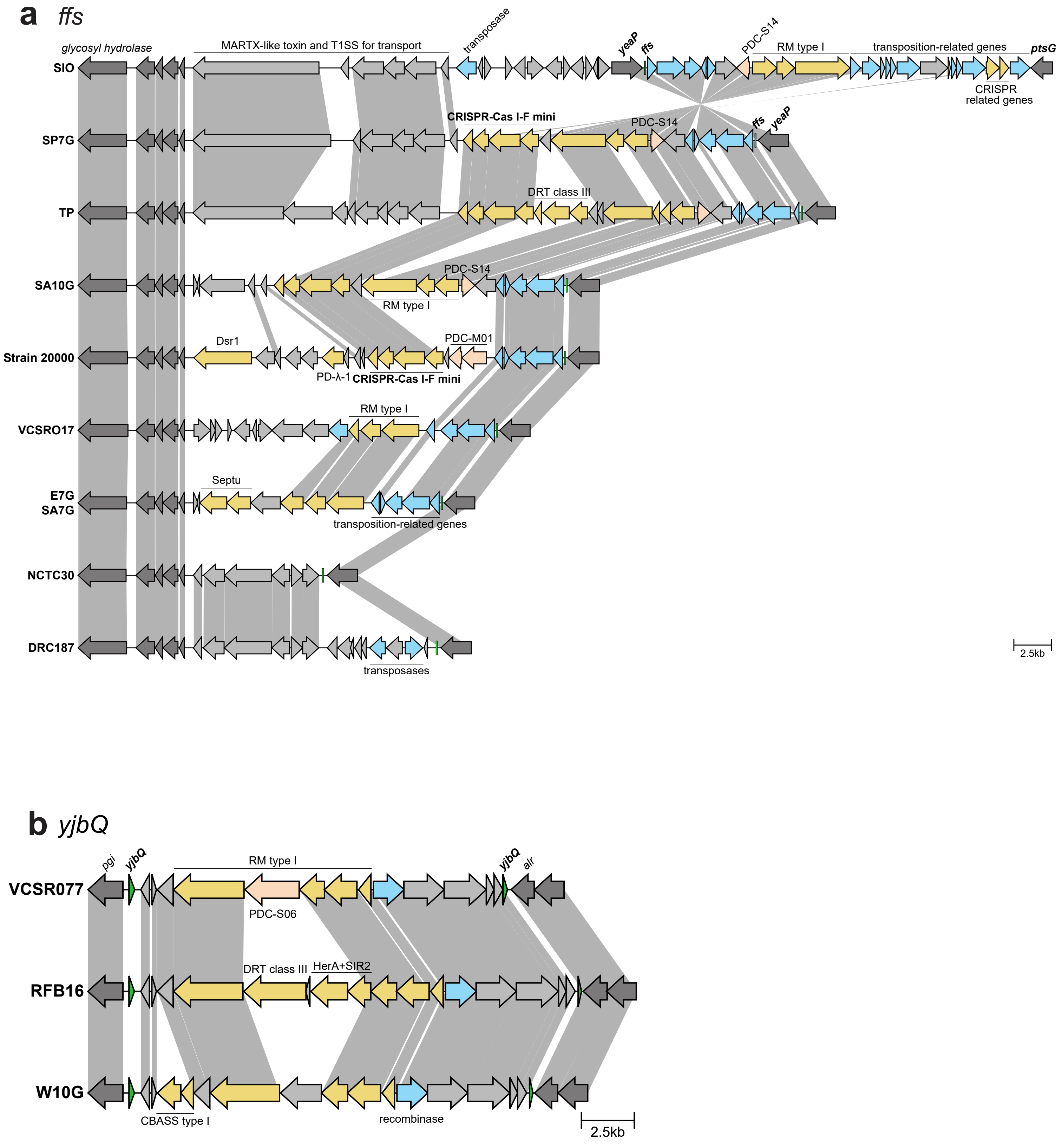

a ffs
glycosyl hydrolase
MARTX-like toxin and T1SS for transport
transposase
yeaP
ffs
PDC-S14
RM type I
transposition-related genes
ptsG
CRISPR related genes
SIO
SP7G
CRISPR-Cas I-F mini
PDC-S14
ffs
yeaP
TP
DRT class III
SA10G
PDC-S14
RM type I
Strain 20000
Dsr1
PDC-M01
PD-λ-1
CRISPR-Cas I-F mini
VCSRO17
RM type I
E7G
SA7G
Septu
transposition-related genes
NCTC30
DRC187
transposases
2.5kb
b yjbQ
pgi
yjbQ
RM type I
yjbQ
alr
VCSR077
PDC-S06
RFB16
DRT class III
HerA+SIR2
W10G
CBASS type I
recombinase
2.5kb

Figure S9

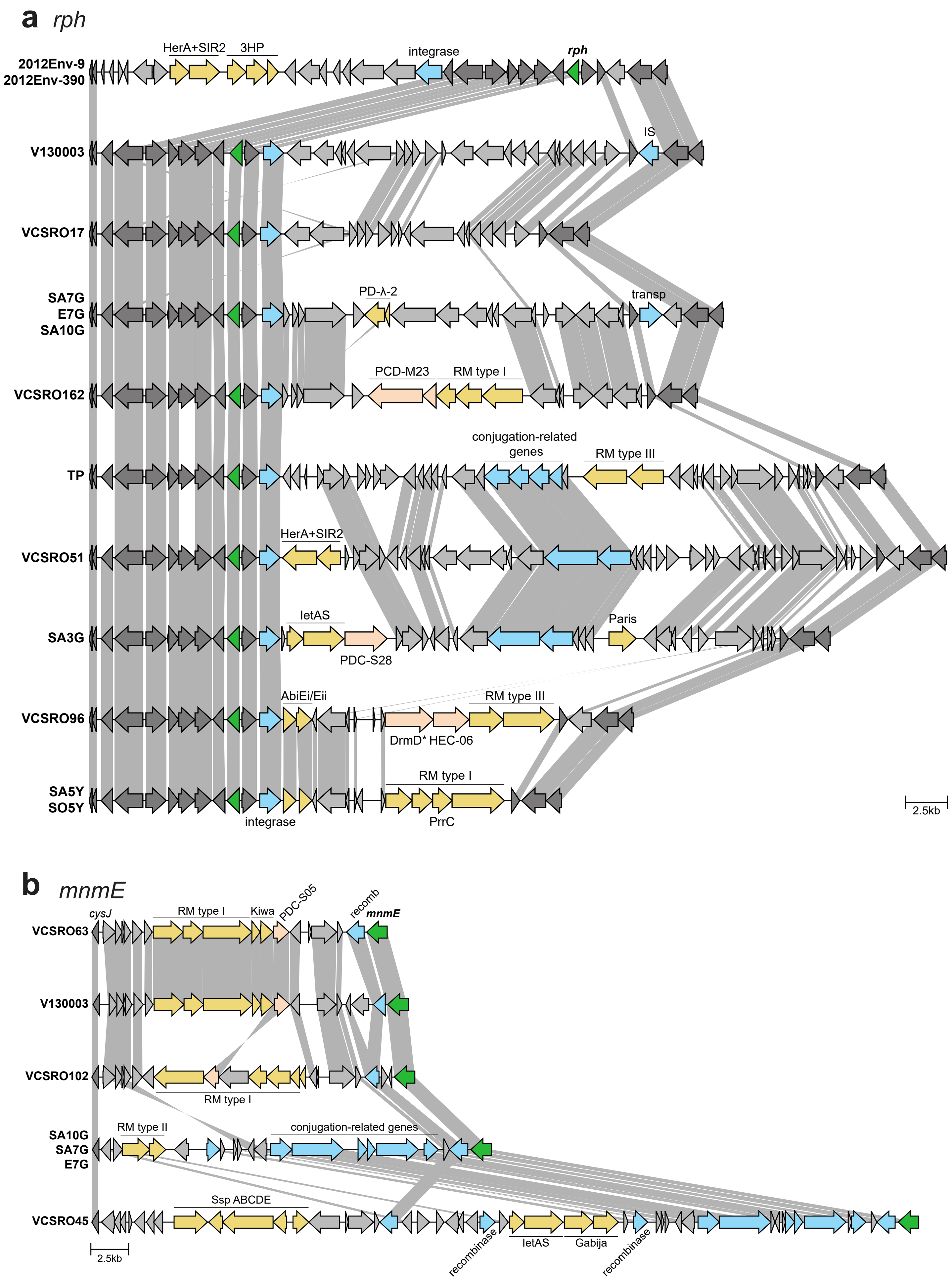
a rph
HerA+SIR2
3HP
integrase
rph
2012Env-9
2012Env-390
V130003
IS
VCSRO17
SA7G
E7G
SA10G
PD-λ-2
transp
VCSRO162
PCD-M23
RM type I
TP
conjugation-related genes
RM type III
VCSRO51
HerA+SIR2
SA3G
letAS
PDC-S28
Paris
VCSRO96
AbiEi/Eii
DrmD* HEC-06
RM type III
SA5Y
SO5Y
RM type I
integrase
PrrC
2.5kb
b mnmE
cysJ
RM type I
Kiwa
PDC-S05
recomb
mnmE
VCSRO63
V130003
VCSRO102
RM type I
SA10G
SA7G
E7G
RM type II
conjugation-related genes
VCSRO45
Ssp ABCDE
recombinase
letAS
Gabija
recombinase
2.5kb

### Figure S10

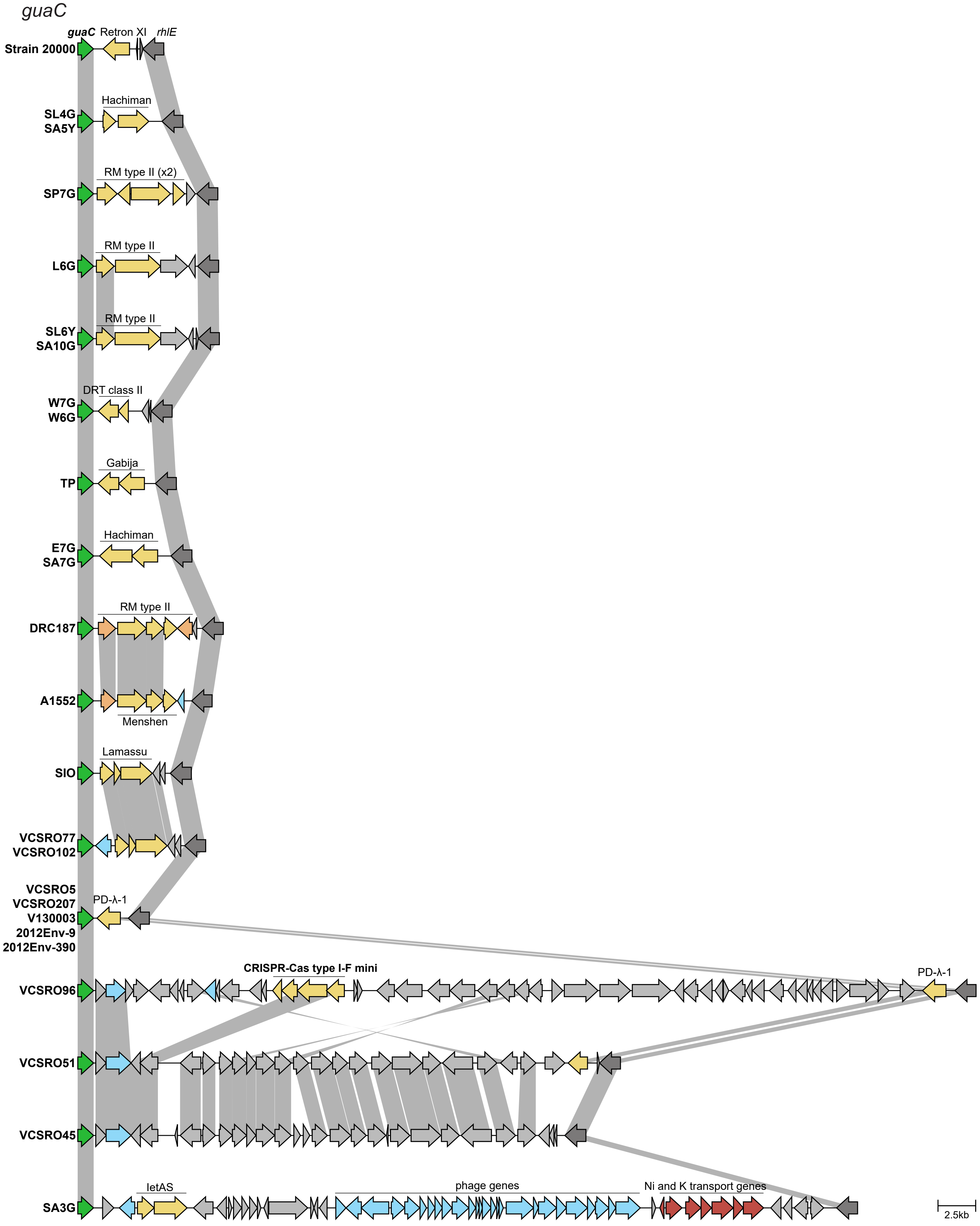
guaC
guaC
Retron XI
rhlE
Strain 20000
Hachiman
SL4G
SA5Y
RM type II (x2)
SP7G
RM type II
L6G
RM type II
SL6Y
SA10G
DRT class II
W7G
W6G
Gabija
TP
Hachiman
E7G
SA7G
RM type II
DRC187
A1552
Menshen
Lamassu
SIO
VCSRO77
VCSRO102
VCSRO5
VCSRO207
V130003
2012Env-9
2012Env-390
PD-λ-1
CRISPR-Cas type I-F mini
PD-λ-1
VCSRO96
VCSRO51
VCSRO45
letAS
phage genes
Ni and K transport genes
SA3G
2.5kb

Figure S11

*VCA0735*

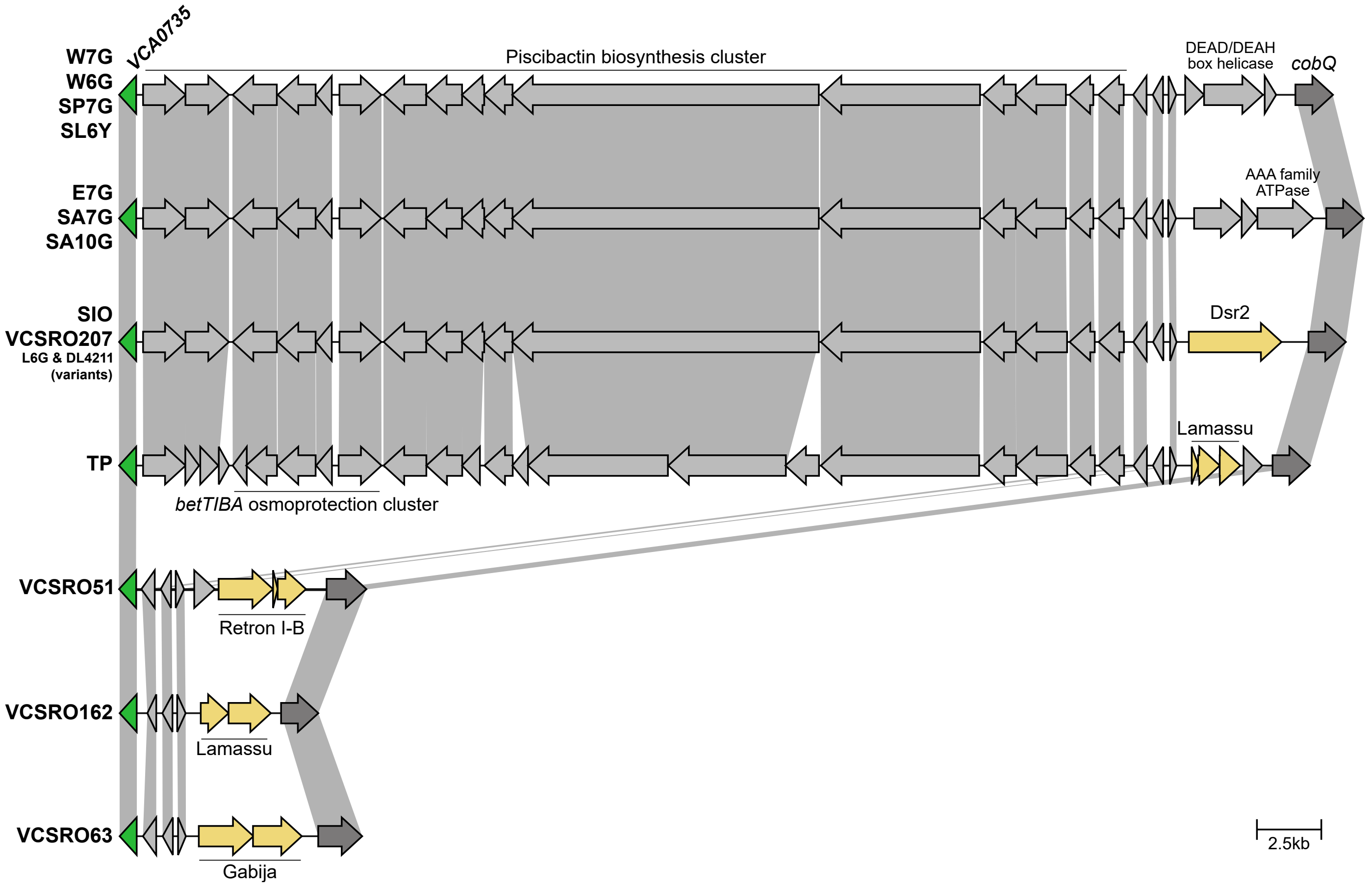

Figure S12

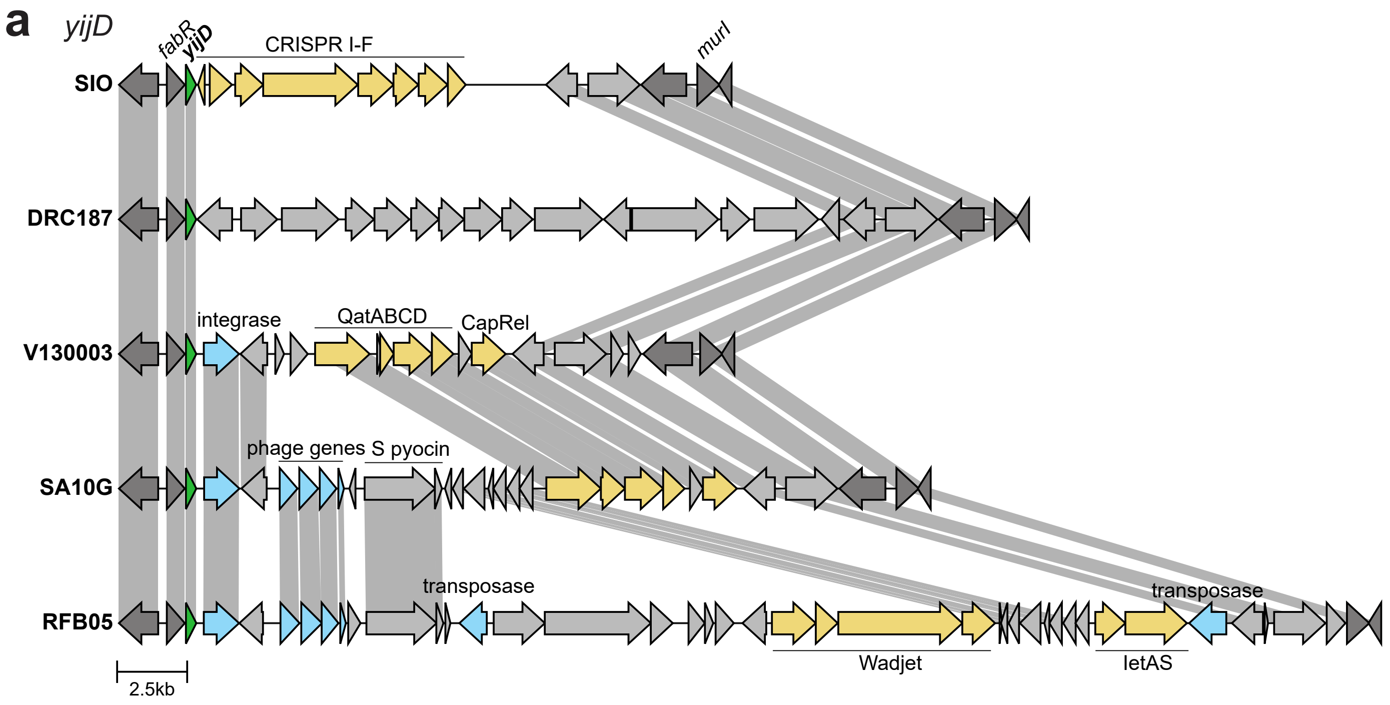
a
yijD
fabR
yijD
CRISPR I-F
murI
SIO
DRC187
integrase
QatABCD
CapRel
V130003
phage genes
S pyocin
SA10G
transposase
transposase
RFB05
Wadjet
letAS
2.5kb

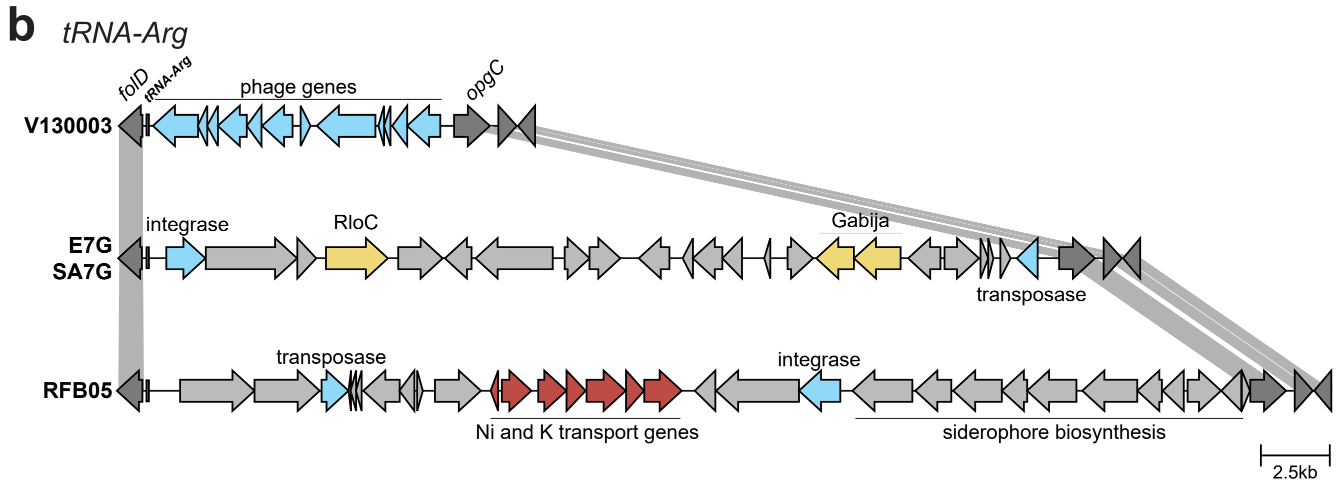
b
tRNA-Arg
folD
tRNA-Arg
phage genes
opgC
V130003
integrase
RloC
Gabija
E7G
SA7G
transposase
transposase
integrase
RFB05
Ni and K transport genes
siderophore biosynthesis
2.5kb
